# A DNA-binding protein tunes septum placement during *Bacillus subtilis* sporulation

**DOI:** 10.1101/459685

**Authors:** Emily E. Brown, Allyssa K. Miller, Inna V. Krieger, Ryan M. Otto, James C. Sacchettini, Jennifer K. Herman

## Abstract

*Bacillus subtilis* is a soil bacterium capable of differentiating into a spore form resistant to desiccation, UV radiation, and heat. Early in spore development the cell possesses two copies of a circular chromosome, anchored to opposite cell poles via DNA proximal to the origin of replication (*oriC*). As sporulation progresses an FtsZ ring (Z-ring) assembles close to one pole and directs septation over one chromosome. The polar division generates two cell compartments with differing chromosomal contents. The smaller “forespore” compartment initially contains only 25–30% of one chromosome and this transient genetic asymmetry is required for differentiation. At the population level, the timely assembly of polar Z-rings and the precise capture of the chromosome in the forespore both require RefZ, a DNA-binding protein synthesized early in sporulation. To mediate precise capture of the chromosome RefZ must bind to specific DNA motifs (*RBMs*) that are localized near the poles around the time of septation, suggesting RefZ binds to the *RBMs* to affect positioning of the septum relative to the chromosome. RefZ’s mechanism of action is unknown, however, cells artificially induced to express RefZ during vegetative growth cannot assemble Z-rings or divide, leading to the hypothesis that RefZ-RBM complexes mediate precise chromosome capture by modulating FtsZ function. To investigate this possibility, we isolated 10 RefZ loss-of-function (rLOF) variants unable to inhibit cell division when expressed during vegetative growth, yet were still capable of binding *RBM*-containing DNA. Sporulating cells expressing the rLOF variants in place of wild-type RefZ phenocopy a Δ*refZ* mutant, suggesting that RefZ mediates chromosome capture through an FtsZ-dependent mechanism. To better understand the molecular basis of RefZ’s activity, the crystal structure of RefZ was solved and wild-type RefZ and the rLOF variants were further characterized. Our data suggest that RefZ’s oligomerization state and specificity for the *RBMs* are critical determinants influencing RefZ’s ability to affect FtsZ dynamics *in vivo*. We propose that RBM-bound RefZ complexes function as a developmentally regulated nucleoid occlusion system for fine-tuning the position of the septum relative to the chromosome during sporulation.

**Author Summary:** The Gram-positive bacterium *B. subtilis* can differentiate into a dormant cell type called a spore. Early in sporulation the cell divides near one pole, generating two compartments: a larger mother cell and a smaller forespore (future spore). Only approximately 30 percent of one chromosome is initially captured in the forespore compartment at the time of division and this genetic asymmetry is critical for sporulation to progress. Precise chromosome capture requires RefZ, a sporulation protein that binds to specific DNA motifs (*RBMs*) positioned at the pole near the site of cell division. How RefZ functions at the molecular level is not fully understood. Here we show that RefZ-*RBM* complexes facilitate chromosome capture by acting through the major cell division protein FtsZ.

## Introduction

To spatially regulate cellular processes, some macromolecules within the cell must assume a hetereogeneous distribution. One way that bacteria create heterogeneity along the bacterial envelope is to utilize proteins that induce and/or partition to sites of membrane curvature^1,2^. From there, membrane curvature proteins can serve as platforms for the localization of additional molecules in the cell. For example, in the rod-shaped bacterium *Bacillus subtilis*, the negative membrane curvature-sensing protein DivIVA coalesces adjacent to past and future cell division sites where it then recruits a cell division regulatory system called Min to inhibit FtsZ polymerization at non-medial sites^3–7^. Another commonly employed mechanism to restrict physiological processes to specific regions of a cell is to require that molecules assemble into larger, multi-subunit complexes to be active. For example, cell division, which requires the coordinated synthesis and turnover of all layers of the cell envelope at midcell, is carried out by a localized multi-subunit complex comprised of over 30 proteins called the “divisome”^8^.

Bacteria also elicit subcellular heterogeneity by harnessing intrinsic properties of macromolecules, such as diffusion rates, oligomerization potential, and affinity for other molecules in the cell^9^. The ParABS system utilized to segregate chromosomes in *Caulobacter crescentus* elegantly demonstrates how bacteria can exploit the intrinsic properties of molecules to achieve spatial regulation. In the ParABS system, proteinprotein and protein-DNA interactions, regulated ATP hydrolysis, and diffusion are harnessed to achieve a new-pole biased gradient of the non-specific DNA-binding ParA. ParB bound at a *parS* site adjacent to *oriC* is initially anchored at the old pole^10,11^. However, once a new round of DNA replication is initiated, affinity of ParB for ParA drives the net movement of the newly formed ParB-*parS* complex (and the replicated *oriC*) toward the ParA-enriched new pole, thus facilitating chromosome segregation^12,13^.

The ParABS system not only demonstrates how the intrinsic properties of molecules can underlie heterogeneity of macromolecules within the cell, but also exemplifies how the nucleoid itself can be utilized in spatial regulation. The nucleoid is highly organized^14^ and many DNA-binding proteins restrict their associated functions to specific cellular addresses by binding to unique DNA motifs. For example, transcription factors only regulate transcription at the promoters they associate with. There are also examples of DNA-binding proteins that bind to specific motifs to regulate the initiation of DNA replication (Spo0J/Soj)^15^, mediate DNA repair and recombination (MutL, XerCD)^16,17^, and segregate chromosomes (ParAB, Spo0J/SMC)^12,13,18,19^. Moreover, some DNA-binding proteins simultaneously interact with the nucleoid and the cell envelope to perform functions in DNA replication (DnaA, SeqA)^20,21^, chromosome organization (RacA, SMC)^22–25^, DNA segregation (FtsK/SpolIIE)^26^, and regulation of cell division (Noc)^27^.

The most extensively studied example of a DNA-binding protein that uses the nucleoid to spatially restrict cell division is SlmA, a TetR family protein found in *Escherichia coli^28^ and several other Gammaproteobacteria including *Vibrio cholerae*^29^. E. coli* SlmA binds to dozens of motifs (SBSs) distributed throughout the chromosome except in the terminus (*ter*) region^30,31^. In a mechanism termed <u>N</u>ucleoid <u>O</u>cclusion (NO), SlmA-SBS complexes inhibit cell division locally by disrupting polymerization of FtsZ^30^,^31^. By restricting SlmA activity to sites of SBS enrichment, *E. coli* effectively inhibits the formation of Z-rings over the bulk nucleoid while at the same time permitting Z-ring assembly in the midcell-localized *ter* region. In this way, SlmA utilizes the chromosome as a landmark to spatially regulate its FtsZ-inhibitory function.

In addition to NO, *E. coli* utilizes at least two other systems to ensure Z-rings only assemble at midcell, between replicated chromosomes. The Min system, alluded to above for *B. subtilis*, inhibits FtsZ polymerization in nucleoid-free regions near the poles^32^. More recently, ZapA, ZapB, and the DNA-binding protein MatP were shown to act in the *ter*-proximal region to promote midcell Z-ring formation^33,34^. Given that reproduction requires the faithful inheritance of intact genomes by progeny, it is not surprising that multiple mechanisms have evolved to ensure chromosomes are faithfully partitioned at the time of cell division.

Like *E. coli*, *B. subtilis* also possesses a NO system to prevent cell division over the bulk nucleoid^35,36^. The NO system of *B. subtilis* is comprised of a DNA-binding protein, Noc, and its cognate binding sites (NBSs), which are also distributed throughout the chromosome except with a notable gap in the *ter* region^36^. In contrast to SlmA, evidence for a direct interaction between Noc and FtsZ is currently lacking. Instead, Noc-NBS complexes associate with the cell envelope, where they are hypothesized to perturb the association and/or nucleation of FtsZ filaments at the membrane^27^.

Establishing and maintaining subcellular organization is important, but cells must also be poised to dynamically reconfigure their overall organization in response to changing growth contexts. For example, during *B. subtilis* sporulation, several major morphological changes must occur to facilitate spore formation. The cell’s two chromosomes are stretched from pole to pole in an elongated *oriC-ter-oriC* configuration called the axial filament^37,38^. In addition, there is a dramatic adjustment in the location of cell division. FtsZ shifts from midcell toward a cell quarter and directs septation over one chromosome. During sporulation, Z-ring inhibition imposed by both the Min and NO systems at the cell pole must be relieved. Alleviation of Min inhibition may be facilitated by the repositioning of MinD (required to mediate MinC-dependent inhibition of FtsZ) to the distal cell pole^39^. Regarding NO, it has been proposed that the axial filament may be arranged such that relatively few Noc-binding sites are positioned at the site of incipient septation^36^.

The shift of FtsZ from midcell toward the pole is promoted by increased levels of FtsZ^40,41^ and expression of a membrane-associated sporulation protein, SpoIIE^42,43^. Following septation, the larger mother cell possesses an entire chromosome, whereas the forespore initially contains only one-quarter to one-third of the second chromosome^18,38^. The genetic asymmetry between the mother cell and forespore is critical for differentiation^44,45^ and the region captured is reproducible^18,38^. The chromosome is not bisected during polar division because SpoIIIE, a DNA translocase localized to the edge of the septum^46^, assembles around the chromosomal arms^26,47^. Since the chromosome is threaded through the septum, SpoIIIE must directionally pump the remainder from the mother cell into the forespore for development to progress. To avoid chromosome breakage during septation, capture a reproducible region of DNA in the forespore, and pump the forespore-destined chromosome in the correct direction, there must be coordination between cell division proteins, SpoIIIE, and the chromosome. How this coordination is orchestrated at the molecular level largely remains a mystery.

Precise division over and capture of the forespore-destined chromosome requires RefZ, a TetR family DNA-binding protein conserved across the *Bacillus* genus^48,49^. RefZ expression is activated early in sporulation, first via the stationary phase sigma factor, *σ*H^50^ and then by Spo0A~P, the activated form of the sporulation master response regulator^51,52^. RefZ binds to five nearly palindromic DNA motifs (RBMs), two on each chromosomal arm and one near *oriC*^48,49^. The *RBMs* on the left and right arms delineate the boundary between chromosomal regions present in the forespore and mother cell at the time of septation. Chromosomal regions immediately adjacent to each *RBM* localize near the incipient site of polar cell division, suggesting a possible role in division or organization of the chromosome near the sporulation septum^48^. Consistent with this idea, the *RBMs* are required for precise capture of the forespore-destined chromosome^48^. Strikingly, the relative position of the *RBMs* with respect to *oriC* is conserved across the entire *Bacillus* genus. This evolutionary conservation strongly suggests that the location of the *RBMs* is functionally important and provides a considerable selective advantage to the genus^48^.

In addition to imprecise chromosome capture, perturbation of RefZ activity is associated with two other phenotypes: first, during sporulation a Δ*refZ* mutant is modestly delayed in assembly of polar Z-rings^49^. Second, artificially induced expression of RefZ during vegetative growth disrupts Z-ring assembly and inhibits cell division. RefZ-DNA complexes are likely required to disrupt Z-rings, as RefZ DNA-binding mutants no longer disrupt cell division^49^. These data, and the fact that RefZ and SlmA are both TetR family proteins led us to hypothesize that *RBM*-bound RefZ complexes might act as a developmentally regulated NO system that tunes FtsZ dynamics and/or Z-ring positioning relative to the chromosome.

To test this hypothesis, we isolated and characterized 10 RefZ loss-of-function (rLOF) variants unable to inhibit cell division when misexpressed during vegetative growth, yet still capable of binding *RBMs*. None of the rLOF variants were able to support wild-type chromosome capture when expressed from the native promoter during sporulation, and instead phenocopied a Δ*refZ* mutant. These results are consistent with a model in which RefZ mediates precise chromosome capture by modulating FtsZ activity. To better understand the molecular basis of RefZ’s activity, wild-type RefZ and the rLOF variants were overexpressed, purified, and structural and biochemical characterizations were carried out. The location of the rLOF substitutions on the RefZ crystal structure suggests that RefZ affects FtsZ through a mechanism that is distinct from that described for SlmA. Characterization of the rLOF variants indicates that specificity for *RBM*-containing DNA and RefZ’s propensity to dimerize are critical determinants in governing RefZ’s effect on cell division and precise capture of forespore chromosome *in vivo*.

## Results

### Identification of RefZ residues important for inhibition of cell division

Misexpression of RefZ during vegetative growth disrupts Z-ring formation and inhibits cell division, resulting in filamentation. The division inhibition phenotype can be suppressed in strain backgrounds harboring specific mutations in *ftsZ* or a second copy of the *ftsAZ* operon^49^. Division inhibition appears to require RefZ’s DNA binding activity, as RefZ variants harboring substitutions in the DNA recognition helix (Y43A and Y44A) do not filament cells following misexpression^49^. DNA binding is also likely required for RefZ’s role in chromosome capture, as a strain harboring point mutations in the five *oriC*-proximal RefZ binding motifs (*RBM_5mu_*) exhibits the same capture defect as a Δ*refZ* mutant^48^. Based on these data, we hypothesized that RefZ associates with *RBMs* to modulate FtsZ dynamics in the vicinity of the incipient septum and that this modulation would be required for ensuring precise chromosome capture.

To test whether RefZ’s ability to inhibit cell division is required to support precise chromosome capture, we designed a two-stage genetic selection-screen to isolate RefZ loss-of-function (rLOF) variants capable of binding to the *RBMs*, but unable to disrupt cell division upon misexpression (Fig 1). Gibson assembly^53^ was used to generate a library of linear misexpression constructs comprised of an IPTG-inducible promoter (P_*hy*_), randomly mutagenized *refZ* sequences (*refZ*\*), a selectable marker (*spec^R^*) and regions of homology to direct double crossover integration of the linear DNA at a nonessential locus (*amyE*)(Fig 1A). To select for rLOF mutants, we took advantage of the fact that in a sensitized background (Δ*minD*), expression of wild-type *refZ* from an IPTG-inducible promoter prevents colony formation on solid medium, whereas expression of RefZ variants unable to inhibit cell division survive^49^. In addition to *minD*, the native *refZ* gene was also deleted to ensure that the only RefZ expressed would be from the inducible promoter.

**Figure 1.**
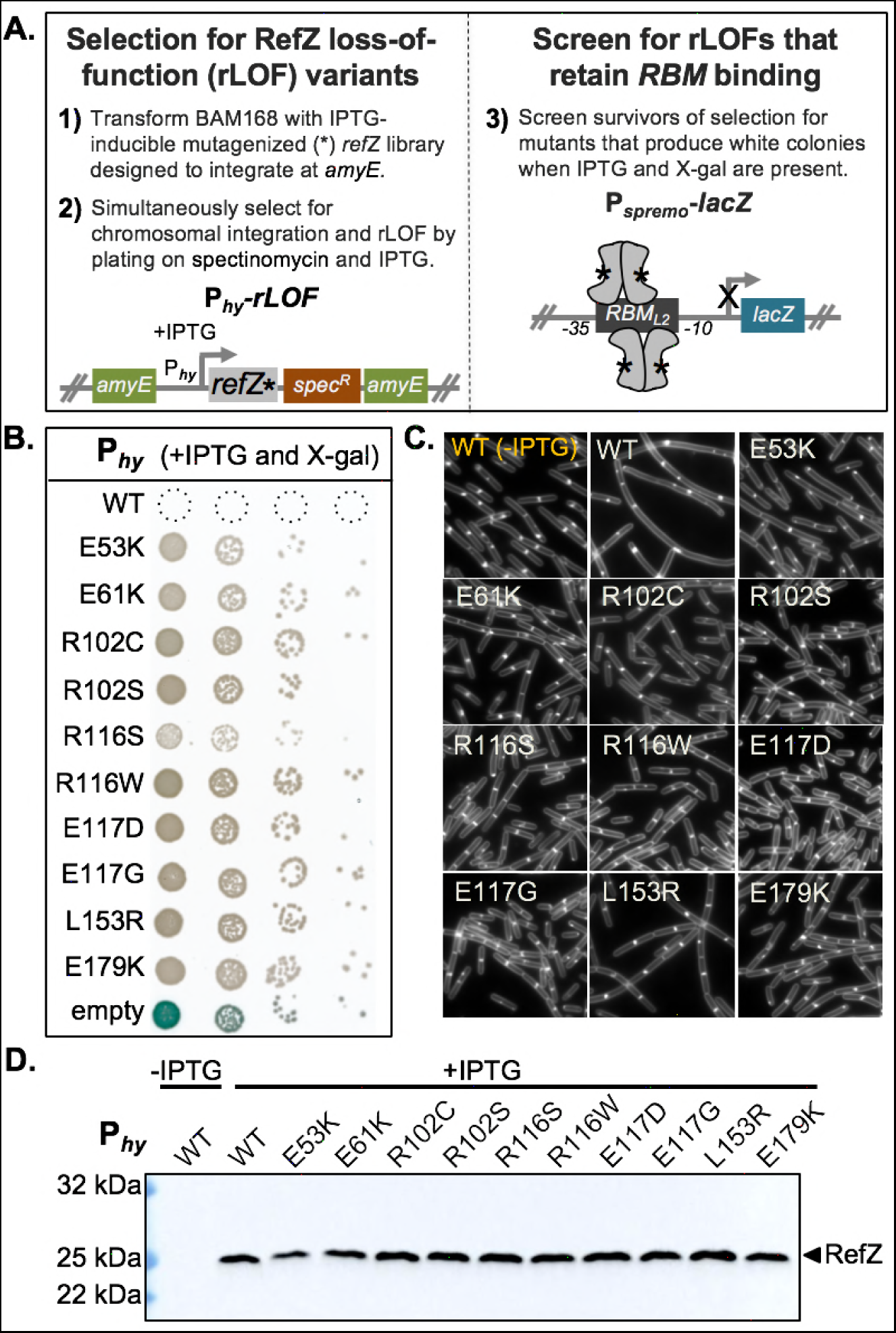
Isolation of rLOF variants. (A) Schematic of genetic selection (left) and screen (right) used to isolate rLOF variants that retain RBM-binding activity. The open-reading frame of *refZ* was mutagenized by error-prone PCR (*refZ*\*), placed under an IPTG-inducible promoter (P_*hy*_), and introduced at the *amyE* locus of competent recipient cells (BAM168). Mutations that interfere with RefZ’s division inhibition function (P_*hy*_-*rLOF*) permit growth in the presence of IPTG. Survivors were screened for *RBM* binding (P_*spremo*_-*lacZ*) on plates containing X-gal and IPTG. (B) Ten unique rLOF variants that do not kill following induction but retain RBM-binding function were identified in the selection-screen. (C) The rLOF misexpression constructs were introduced into a wild-type (*Bs168*) genetic background and the extent of cell filamentation in CH medium following 90 min of induction with 1 mM IPTG was monitored using epifluorescence microscopy. Membranes were stained with TMA (white). The uninduced wild-type (WT) control is labeled in yellow. (D) Western blot analysis to monitor the production and stability of wild-type RefZ (WT) and the rLOF variants following 45 min of induction with 1 mM IPTG. RefZ is not produced at levels detectable above background with our antibody during vegetative growth (Lane 1, uninduced) or sporulation (data not shown).

To eliminate variants unable to bind DNA, survivors of the selection were screened for RBM-binding activity using a RefZ-repressible, *lacZ* transcriptional fusion (P*_spremo_-lacZ*) integrated at the non-essential sacA locus. P_*spremo*_ harbors a single *RBM* (*RBM_L2_*)^48^ inserted between the −35 and −10 elements of a constitutive promoter (Fig 1A). In this background, rLOF variants that can bind the engineered *RBM* operator repress *lacZ* expression and produce white colonies on media containing X-gal. In contrast, rLOF variants unable to bind the *RBM* due to decreased affinity for the *RBM*, poor expression, truncation, or misfolding produce blue colonies, allowing them to be excluded from further investigation.

To facilitate selection and screening efficiency and avoid cloning steps, transformation conditions were optimized so that the mutant *refZ* misexpression construct library could be directly introduced into the *B. subtilis* chromosome (see Methods). RefZ loss-of-function and double-crossover integration were selected for simultaneously by plating transformations on a medium containing both spectinomycin and IPTG.

Approximately 1,300 viable transformants were obtained, 37 of which were either white or pale blue on medium containing X-gal and IPTG, consistent with rLOF repression of *lacZ* expression from the engineered *RBM* operator. Since resistance to RefZ can also be conferred by spontaneous suppressor mutations in *ftsZ*^49^, the 37 misexpression constructs were transformed into a clean selection-screen background, and survival and RBM-binding were reassessed. Four candidates failed to survive on IPTG plates, suggesting the presence of suppressor mutations in the original strains, while an additional four turned blue on X-gal indicator medium.

To identify rLOF mutations in the remaining 29 candidates, the P_*hy*_-*rLOF* region was amplified from the genomic DNA and sequenced. Ten unique single-point mutations were identified, corresponding to the 10 rLOF substitutions shown in Figure 1B. In contrast to wild-type RefZ, misexpression of the rLOF variants did not result in cell filamentation (Fig 1C), consistent with a loss of ability to affect FtsZ. The inability of rLOF variants to inhibit cell division was not anticipated to be attributable to protein misfolding or insufficient expression, as each variant was able to repress *lacZ* expression from the *RBM* operator in the primary screen (Fig 1B). Consistent with this conclusion, Western blot analysis of the rLOF variants demonstrated that they are stably expressed and present at levels comparable to wild-type RefZ following misexpression (Fig 1D). From these data we conclude that the 10 rLOF variants are perturbed in their ability to affect FtsZ function, either directly or indirectly.

### rLOF mutants miscapture the forespore chromosome

A Δ*refZ* mutant and a strain harboring point mutations in all five ori’C-proximal *RBMs* (*RBM_5mu_*) both exhibit a 2-fold increase in the frequency of left and right arm reporter capture compared to wild-type controls^48^. We hypothesized that if RefZ’s ability to perturb FtsZ assembly is required to mediate precise chromosome capture, then the rLOF mutants would phenocopy the Δ*refZ* mutant with regard to chromosome trapping. To test this hypothesis, chromosome organization was monitored in sporulating cells expressing the rLOF variants from the native locus (native promoter) using a fluorescence-based trapping assay^18,48^. For each strain, the native *refZ* gene was replaced with a rLOF mutant sequence in backgrounds harboring reporters for either left (−61°) or right (+51°) arm capture (Fig 2). All of the rLOF mutations resulted in significant increases in both left and right arm reporter capture compared to wild-type controls (P<0.05)(Fig 2). Moreover, with the exception of right arm capture in the R116S mutant, miscapture of both left and right arm reporters in the rLOF mutants was statistically indistinguishable from the Δ*refZ* controls (P>0.05). The right arm reporter in the R116S mutant exhibited an intermediate capture defect that was statistically different from both Δ*refZ* (P=3.9×10^−3^) and wild-type (P=2.3×10^−3^). The intermediate capture defect observed in the R116S mutant suggests this variant retains some functionality, and is consistent with the reduced growth we observed on selection medium in the sensitized Δ*minD* background (Fig 1B). These data demonstrate that the same residues required for RefZ’s ability to inhibit division upon misexpression are also required for precise chromosome capture, and are consistent with a model in which *RBM*-bound RefZ modulates FtsZ activity to position the polar septum relative to the chromosome.

**Figure 2.**
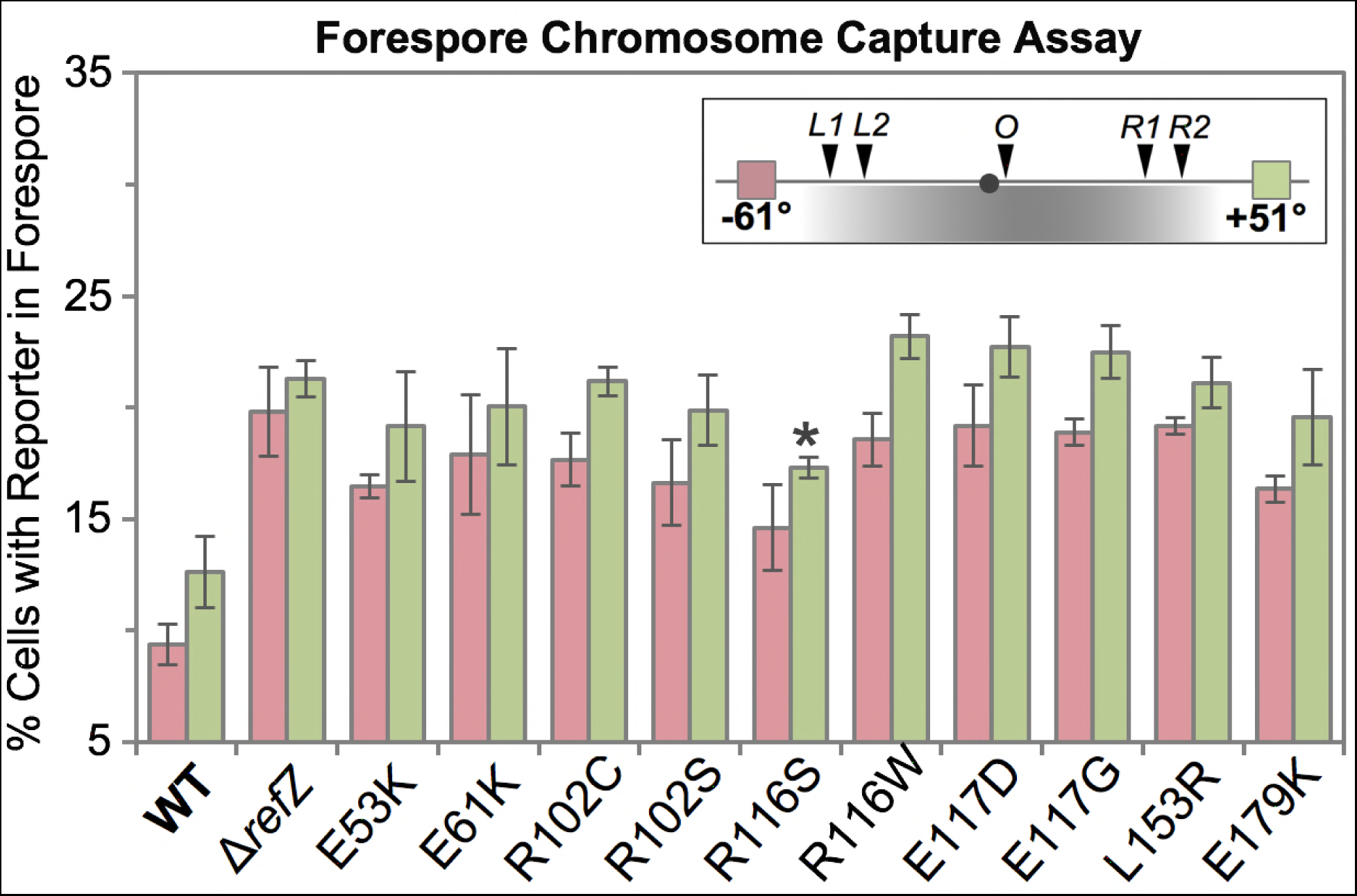
rLOF variants unable to inhibit cell division miscapture regions of the forespore chromosome. Quantitative single cell analysis of chromosome capture is represented as the average percentage of cells that captured either the left arm (−61°, pink) or right arm reporter (+51°, green) in the forespore at the time of polar division. The black circle represents *oriC* (0°). The inset indicates the location of the reporters relative to the *RBMs*, with the region of chromosome typically captured in the forespore shaded grey. All strains encoding rLOF variants miscapture the left and right arm reporters at levels statistically indistinguishable from the Δ*refZ* mutant control (P>0.05) with the exception of the R116S variant. The R116S right arm reporter exhibited an intermediate capture defect that was statistically different from both Δ*refZ* (asterisk, P=3.9×10^−3^) and wild type (P=2.3×10^−3^). Error bars represent standard deviations.

### Structural characterization of RefZ

Like the *E. coli* NO protein, SlmA, RefZ belongs to the TetR family of DNA-binding proteins^49^. At the sequence level, RefZ and SlmA share no significant similarity. We reasoned that structural characterization of RefZ and mapping of the rLOF substitutions to the RefZ structure would not only provide insight into how RefZ functions, but also allow for comparison to what is known about SlmA’s mechanism of FtsZ inhibition. RefZ-His6 was purified, crystallized, and the structure was solved using single-wavelength anomalous dispersion (SAD) phasing at a resolution of 2.6 Å. RefZ crystallized as a homodimer (Fig 3A) with one molecule in the asymmetric unit of a P4_1_2_1_2 crystal lattice. The model for residues 1-200 was built and refined with R_work_= 22% and R_free_= 25% (Table 1). Each RefZ subunit is composed of 10 α-helices connected by loops and turns, with α1, α2, and α3 comprising the DNA binding helix-turn-helix (HTH) domain and α4-α10 comprising the regulatory domain (Fig 3A), similar to other structurally characterized TetR family proteins^54^. There are two major regions for dimerization contacts. Helices α7, α8, α9, and α10 form regulatory domain contacts with α7’, α8’, α9’, and α10’, with α8, α10, α8’ and α10’ forming a four-helix dimerization motif (Fig 3B). A second interface is formed by α6 and α6’, at the junction between the regulatory and DNA binding domains (Fig 3A). Although the crystallization condition included *RBM*-containing DNA, we observed no DNA in the crystal structure. In fact, the HTH DNA binding domain is involved in extensive crystal packing interactions, likely precluding DNA binding within the crystal lattice.

**Figure 3.**
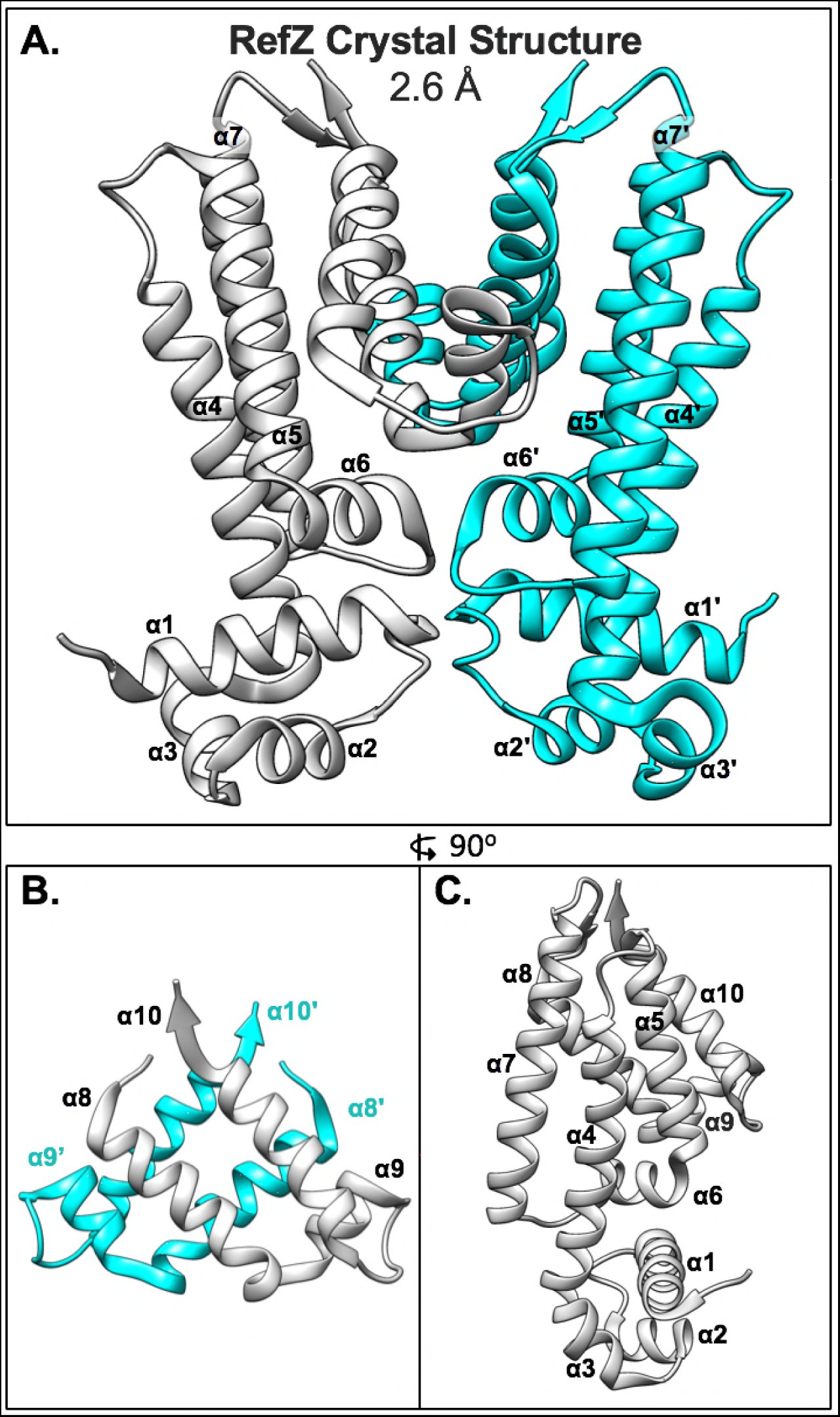
Crystal structure of the RefZ homodimer at 2.6 Å resolution. (A) Structure of the RefZ homodimer. Subunits are colored grey and cyan. (B) Helices α8-α10 of RefZ’s regulatory region with antiparallel helices α8, α10, α8’, and α10’ comprising the four-helix dimerization motif. (C) The RefZ monomer, rotated 90° relative to panel A.

**Table 1.** Data collection, phasing and refinement statistics for the RefZ structure.

| PDB ID | 6MJ1 |
| --- | --- |
| **Data collection** |  |
| Space group | P 4<sub>1</sub> 2<sub>1</sub> 2 |
| Cell dimensions |  |
| *a*, *b*, *c* (Å) | 100.021, 100.021, 100.177 |
| $\alpha$, $\beta$, $\gamma$ (°) | 90, 90, 90 |
| Resolution (Å) | 2.6 |
| *R*<sub>merge</sub> | 0.11 (0.79) |
| *I* / σ*I* | 11.59 |
| Completeness (%) | 100 (100) |
| Redundancy | 17.6 (15.6) |
| **Refinement** |  |
| Resolution (Å) | 44.952-2.6 |
| No. reflections | 16,039 |
| *R*<sub>work</sub> / *R*<sub>free</sub> | 22.20 / 25.36 |
| No. atoms |  |
| Protein | 1,683 |
| Water | 25 |
| *B*-factors |  |
| Protein | 76 |
| R.m.s. deviations |  |
| Bond lengths (Å) | 0.009 |
| Bond angles (°) | 1.082 |

According to a structural similarity search using VAST^55^, RefZ shares the highest homology with PfmR from *Thermus thermophilus* (PDB: 3VPR)^56^, with a VAST similarity score of 15.4, closely followed by KstR2 of *Mycobacterium tuberculosis* (PDB: 4W97)^57^, with a score 15.2. The SlmA structure (PDB: 4GCT)^58^ was the tenth closest in similarity with a score of 13.6. Superposition of SlmA and RefZ produced a root-mean-square deviation (rmsd) in Cα of 2.8.

RefZ’s HTH domain (residues 1-45) has the highest contiguous alignment similarity score with QacR from *Staphylococcus aureus* (PDB: 1JT6)^59^, with a VAST similarity score of 4.0 and a rmsd value of 0.7. Superimposition of the HTH domains demonstrates the structures align closely (S1A Fig). However, when the RefZ dimer is superimposed with DNA-bound QacR (PDB: 1JT0), it is apparent that the RefZ dimer would need to undergo a conformational change for the α3 and α3’ helices to be accommodated in adjacent DNA major grooves (S1B Fig and S1C Fig).

DNA binding in TetR family proteins can be allosterically regulated by ligand binding in a pocket formed by α5, α6, and α7. For QacR, ligand binding results in a pendulum motion of α4 that repositions the HTH domains such that the distance between α3 and α3’ becomes incompatible with DNA binding^60^. In the RefZ structure (unbound from DNA), there is no obvious ligand binding pocket in the α5-α7 regulatory region, suggesting RefZ’s affinity to DNA is unlikely to be regulated by ligand binding in this region. At the same time, we do not exclude the possibility that a pocket may exist when RefZ is bound to DNA.

### The regions of RefZ and SlmA important for inhibiting cell division are distinct

To analyze which regions of RefZ are important for its effect on cell division, and compare them to the location of the loss-of-function residues identified for SlmA, the residues with rLOF substitutions were mapped to the RefZ crystal structure (Fig 4).

**Figure 4.**
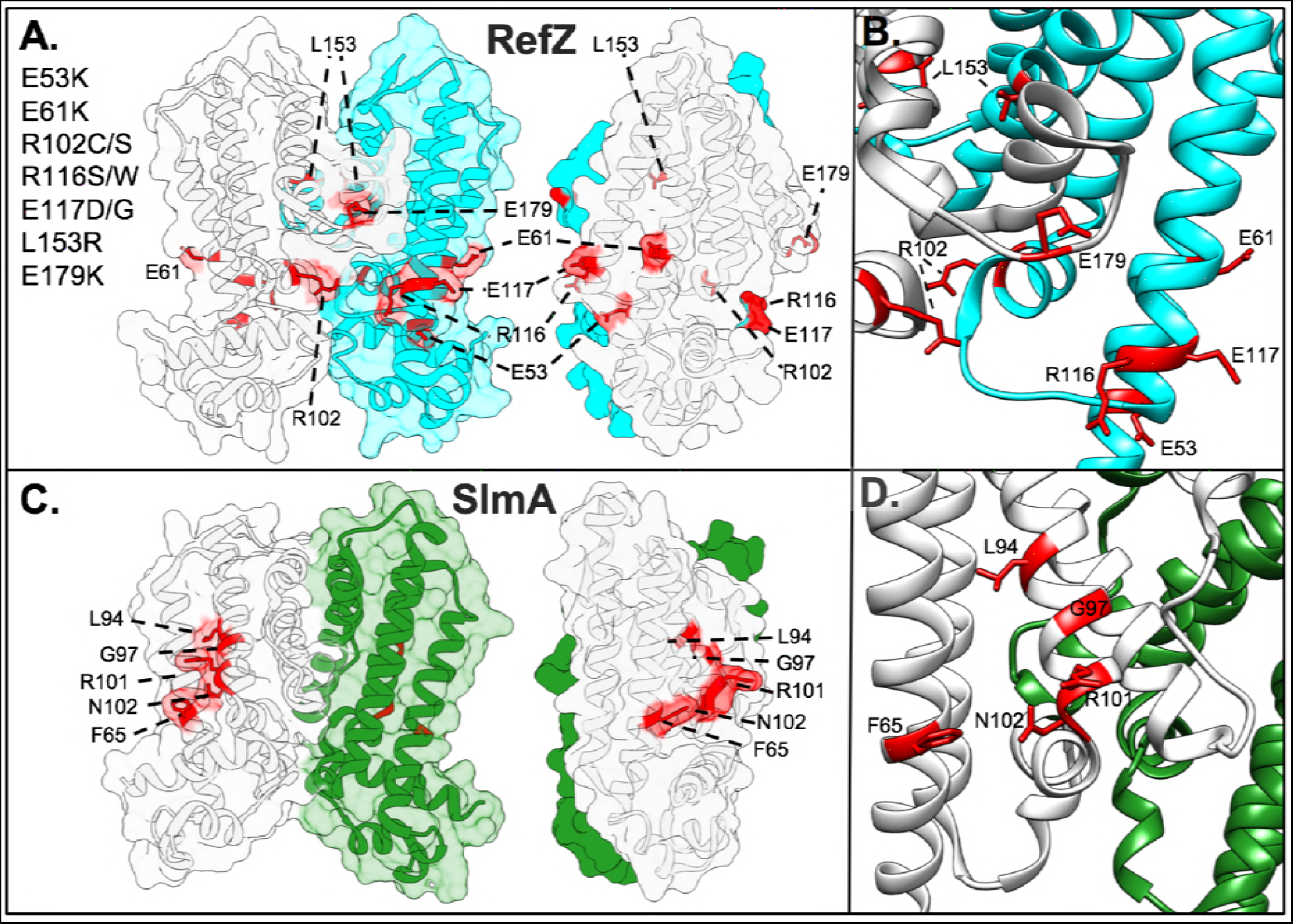
Position of residues implicated in RefZ’s regulation of cell division. (A) Surface/cartoon representation of the RefZ homodimer highlighting residues with substitutions conferring loss-of-function (red, sticks). Subunits are colored white and cyan. (B) Ribbon model of RefZ region showing residues conferring loss of function as sticks. (C) Surface/cartoon representation of the SlmA homodimer (PDB: 5HBU) highlighting residues with substitutions conferring loss of function (red, sticks). Subunits are colored white and green. (D) Ribbon model of SlmA region showing residues conferring loss of function as sticks.

Nine of the 10 rLOF substitutions (L153R being the exception) occur in charged residues that are surface exposed and map to the same surface of the RefZ homodimer (Fig 4A and 4B). L153 maps to the dimerization interface (Fig 5A) and participates in several hydrophobic interactions between subunits that are likely important for RefZ dimerization. Residue R102 is not only surface exposed, but also hydrogen bonds across the dimer interface to the backbone carbonyl of V108’ (NH_2_-O = 2.6 Å)(Fig 5B).

**Figure 5.**
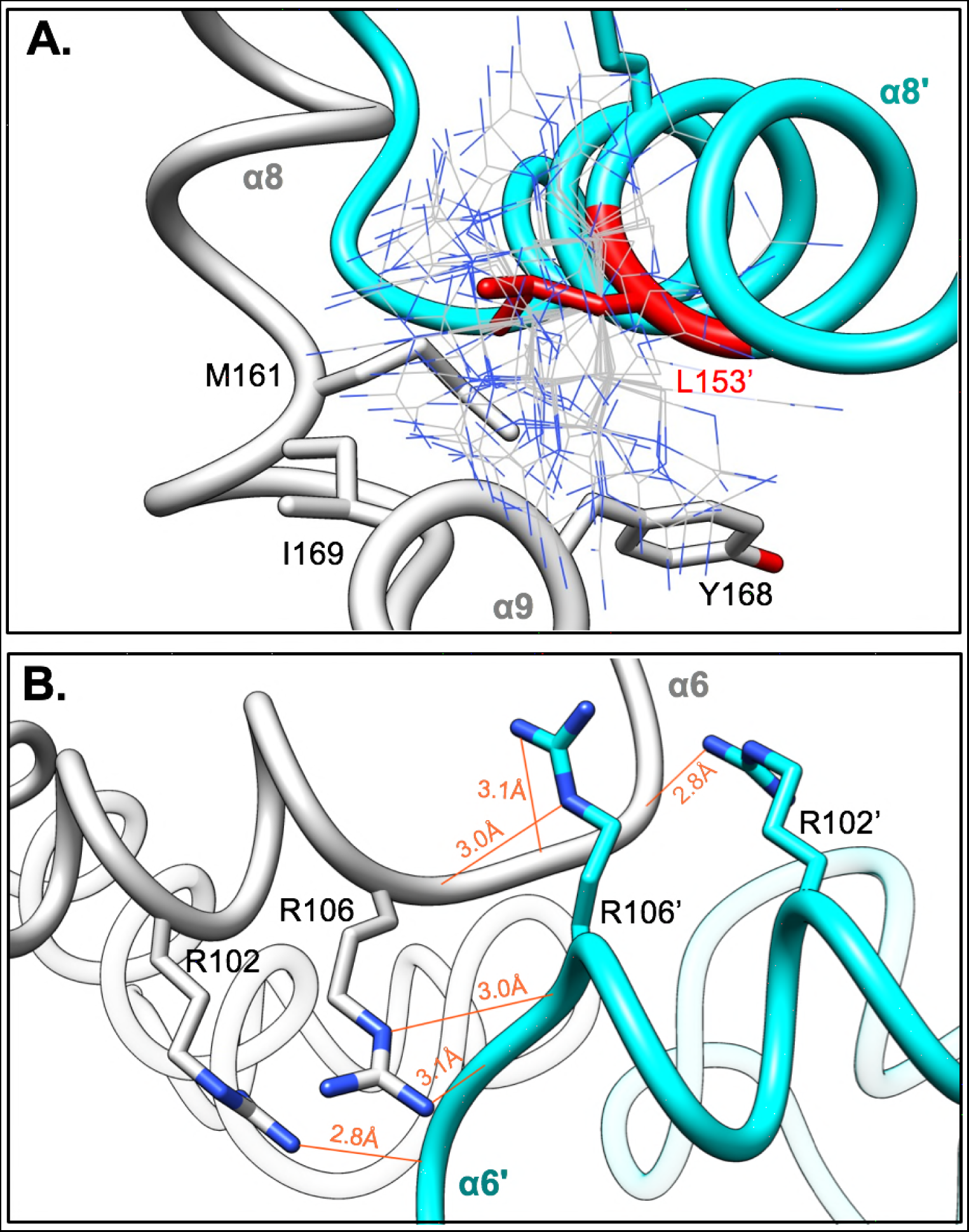
Dimer interface residues implicated in RefZ function. RefZ subunits are shown in light gray and cyan. (A) Hydrophobic dimerization interface near the L153 residue. Thin blue and gray sticks display possible positions of an R153 side-chain based on a rotamer library. (B) Helices α6 and α6’ of RefZ with residues implicated in loss of function shown as sticks. The hydrogen bonds formed across the dimer interface by R102 and R106 are displayed as red lines.

To assess if similar regions of SlmA were implicated in FtsZ regulation, the structures of the RefZ and SlmA homodimers were compared (Fig 4). SlmA binds the C-terminal domain tail of FtsZ along a hydrophobic groove located between α4 and α5^29,30^. SlmA loss-of-function substitutions map to this region clustering primarily along α4 (Fig 4C and 4D)^29,61^. In contrast, the surface-exposed residues implicated in RefZ loss of function are positioned both at or on either side of the RefZ dimerization interface and all but L153 are positively or negatively charged (Fig 4A). From these data we conclude that if RefZ regulates FtsZ through direct interaction, then the precise mechanism is likely to differ significantly from that described for SlmA.

### Characterization of RefZ and rLOF variant DNA-binding

RefZ’s ability to inhibit cell division is dependent upon DNA binding^49^. We predicted that the rLOF variants would be DNA-binding proficient because each was able to repress *lacZ* expression from an *RBM* operator in the *in vivo* screening assay (Fig 1B); however, RBM-binding in the *in vivo* assay was qualitative and not designed to differentiate between specific and non-specific DNA interactions. To directly examine the behavior of the variants with DNA, we overexpressed and purified each of the rLOF variants (S2 Fig) and performed electrophoretic mobility shift assays (EMSAs) with wild-type and mutant *RBM* DNA probes as described previously^48^. Incubation of wild-type RefZ with a 150 bp *RBM*-containing probe produced two major mobility shifts (Fig 6), corresponding to RefZ binding to *RBM*-containing DNA in units of two and four. Consistent with previous observations^48^, the upshifts were lost when RefZ was incubated with a mutant *RBM* probe (harboring seven point-mutations in the central palindrome) suggesting that DNA binding is shows specificity for the *RBM* sequence (Fig 6). Four of the rLOF variants (R116S, R116W, E117D, and E179K) produced specific upshifts similar to wild-type RefZ, suggesting that their loss-of-function phenotypes are not attributable to altered affinity or non-specific DNA binding.

**Figure 6.**
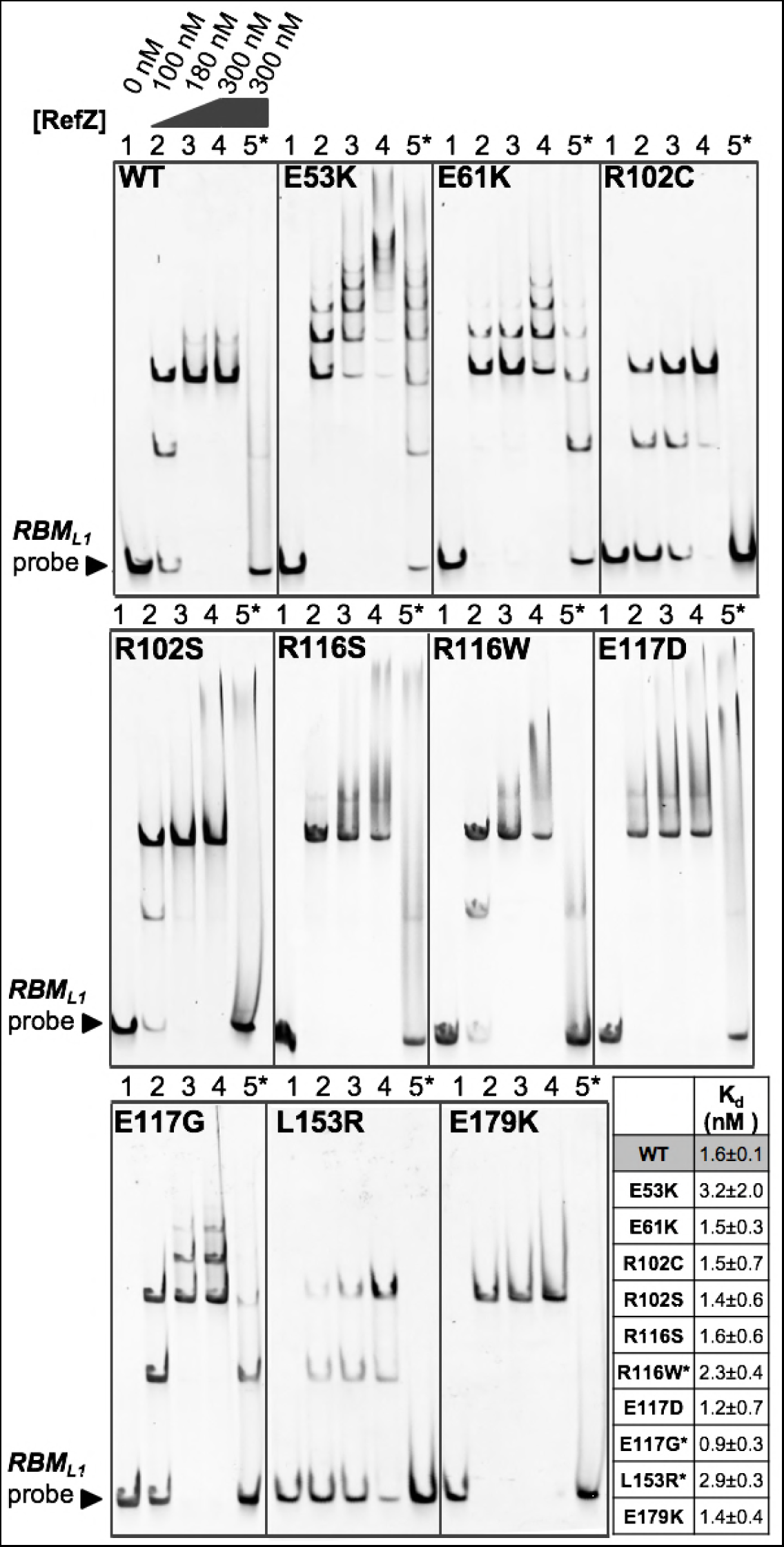
Interaction of the rLOF variants with DNA. Electrophoretic mobility shift assays were performed with 150 bp DNA probes (10 nM) centered on either the wild-type (lanes 1-4) or the mutant (lane 5*) *RBM_L1_* sequence. Probes were incubated with the indicated concentrations of purified RefZ-His6 (WT) or rLOF-His6 variants for 30 min. Reactions were run on a 5% TBE gel for 30 min at 150 V. The tabulated K_d_ values of RefZ for an immobilized 41 bp *RBM*-containing DNA segment were determined using a bio-layer interferometry assay. All the variants possessed K_d_ values within 2-fold of the wild-type K_d_. The differences in K_d_ between wild-type RefZ and R116W, E117G, and L153R are significant (indicated by asterisks)(P=0.05, P=0.025, and P=0.003, respectively).

The remaining variants exhibited altered DNA interactions with respect to either specificity and/or mobility shift pattern. Two variants (E53K and E61K) exhibited a laddering pattern, possibly due to additional subunits of RefZ binding nonspecifically along the DNA (Fig 6). These variants also shifted a mutant *RBM*, consistent with enhanced nonspecific binding. E53K and E61K may assume conformations more favorable for nonspecific DNA binding since the substitutions are located on α4, a helix important for modulating DNA interaction in response to ligand binding in other TetR family members^62^. Although the laddering behavior was most extensive with E53K and E61K mutants, wild-type RefZ is also observed to ladder slightly (Fig 6). The laddering behavior is more apparent when the EMSA gels are run at a higher voltage (200 V vs. 150 V)(S3A Fig), likely because EMSAs are non-equilibrium assays and the faster run time reduces RefZ disassociation. E117G also produced laddering, albeit to a lesser extent than either E53K or E61K (Fig 6). The remaining variants, R102C, R102S, and L153R, each possess substitutions in residues that make dimerization contacts (Fig 5). R102C, R102S and L153R produced two major upshifts, but were unable to ladder on DNA even under EMSA conditions in which wild-type RefZ displayed some laddering (S3B Fig).

To determine if there were quantitative differences in DNA binding that might account for the loss-of-function phenotypes, we determined the dissociation constant (K_d_) of wild-type RefZ and each of the rLOF mutants for a 41 bp segment of *RBM*-containing DNA using bio-layer interferometry. The *RBM*-containing DNA, which was 5’ biotinylated, was immobilized on a streptavidin sensor. The association and dissociation of wild-type RefZ (S3 Fig) and the rLOF variants was then assessed by monitoring the change in thickness of the bio-layer. All of the rLOF variants displayed K_d_ values within 2-fold of wild type (Fig 6). The decreased K_d_ for the L153R mutant was most significant (P<0.01), consistent with the reduced apparent affinity for DNA observed by EMSA (Fig 6). These results suggest that the *in vivo* chromosome capture defect observed in strains harboring rLOF mutations (Fig 2), with the possible exception of L153R, are unlikely attributable to markedly reduced affinity for DNA.

### RefZ oligomerization state by size-exclusion chromatography

Three of the rLOF substitutions (R102C, R102S, and L153R) map to residues implicated in RefZ dimerization based on structural analysis (Fig 5), suggesting dimerization may be important for RefZ’s effect on cell division. Purified TetR proteins have been shown to exist as both monomers and dimers in solution and as pairs of dimers on DNA^31,62–65^. RefZ also binds DNA in units of two and four^48^, but its oligomerization state in the absence of DNA is unknown. To determine the oligomerization state of purified RefZ and the rLOF variants, we performed size-exclusion chromatography. Wild-type RefZ-His6 eluted from a Superdex 200 column primarily as a single peak corresponding to an apparent molecular weight of 21 kDa, close to the actual monomeric molecular weight of 25.4 kDa (Fig 7A and S4 Fig). A minor peak, corresponding to an aggregate or higher-order oligomer, was also observed (S4 Fig). All of the rLOF variants tested displayed elution profiles comparable to wild type (Fig 7A). These data indicate that if RefZ forms dimers in the absence of DNA, then these dimers are not stable enough to be maintained during size-exclusion chromatography.

**Figure 7.**
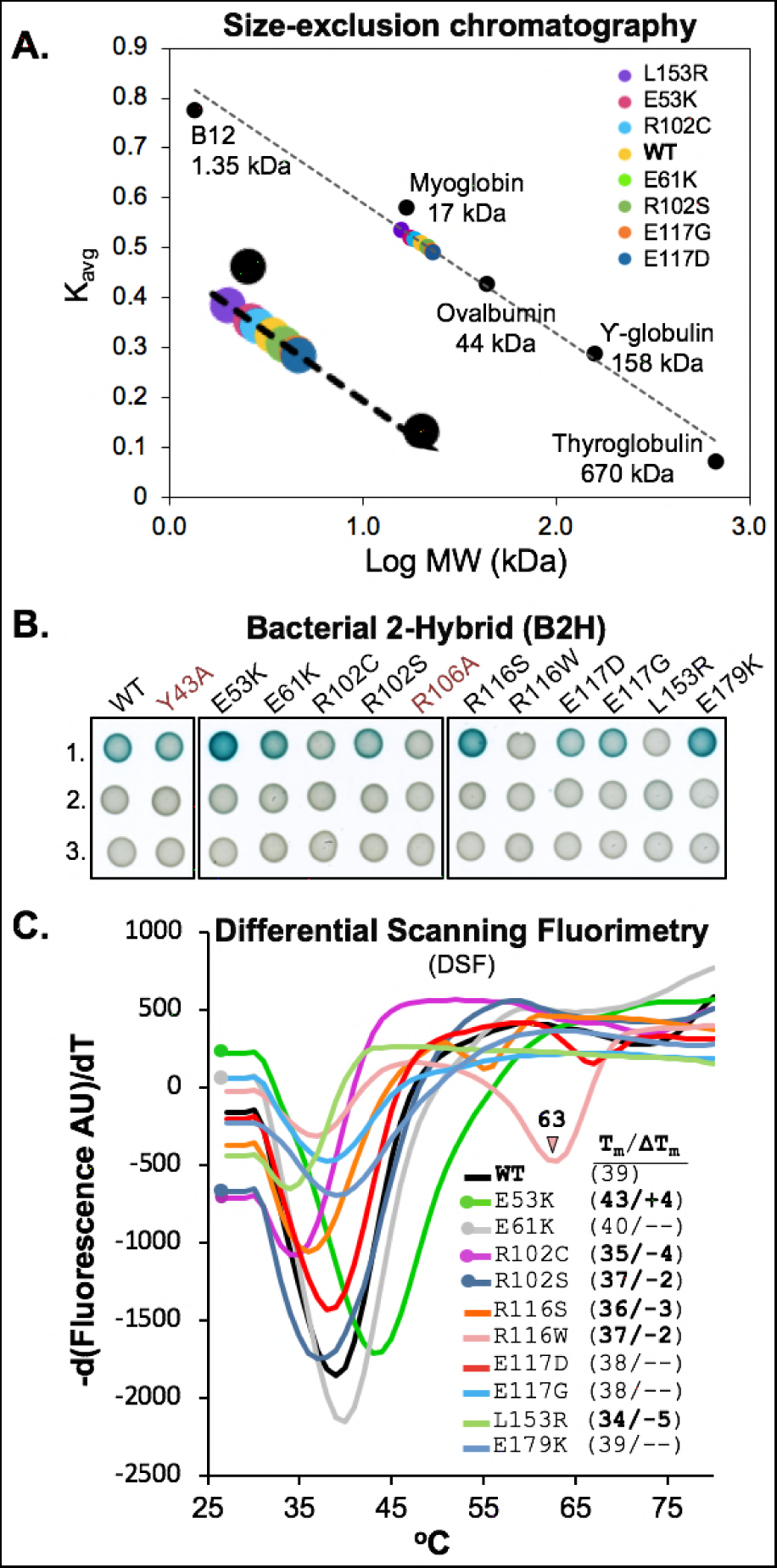
Oligomeric state and thermostability of wild-type RefZ and the rLOF variants. (A) Size-exclusion chromatography of wild-type RefZ-His6 and a subset of rLOF-His6 variants on a Superdex 200 column. The K_avg_ values for the indicated standards were used to generate a standard curve and to estimate the apparent molecular weights of the experimental samples. The E61K and R102C variants share the same position on the curve and only R102C (cyan) is visible. (B) Self-interaction of wild-type RefZ or rLOF variants in a B2H assay. The RefZ variants in red (Y43A and R106A) were generated by site-directed mutagenesis and do not bind *RBM*-containing DNA. Wild-type RefZ subunits or the subunits of the indicated variants were fused to T25 and T18 tags. Pairwise interactions between wild-type RefZ subunits or the subunits of the indicated variants fused to T25 and T18 tags (row 1), T25 tagged subunits paired with an empty T18 vector (row 2), or T18 tagged subunits paired with an empty T25 vector (row 3). Color development after 41 h of growth at room temperature is shown. (C) DSF of wild-type RefZ-His6 and the rLOF-His6 variants. Protein stability is reported by fluorescence of SYPRO orange as a function of increasing temperature. T_m_ values were calculated by determining the temperature at which the first derivative, d(Fluorescence AU)/dT, is at a minimum. ΔT_m_ (inset) is the difference in T_m_ values between wild-type RefZ and each rLOF variant. A ΔT_m_ value of 1.5°C or less was not considered to be significant, and is shown as a dash.

### Bacterial two-hybrid analysis of RefZ self-interaction

Size-exclusion chromatography is known to disassociate weaker oligomers, including dimers of at least one TetR family protein^60^. Therefore, to further investigate if any of the rLOF substitutions altered RefZ’s ability to form dimers, we performed bacterial 2-hybrid (B2H) analysis^66^. In the B2H assay, wild-type RefZ displayed a self-interaction that was not observed in the negative controls (Fig 7B). The self-interaction is unlikely to require *RBM* binding, as the B2H assay is performed in an *E. coli* strain that lacks native *RBM* motifs. Consistent with this observation, a DNA-binding deficient variant, Y43A^49^, displayed self-interaction similar to wild type (Fig 7B). The B2H is most likely reporting on dimerization as the RefZ forms a homodimer in the crystal structure (Fig 3A). To explore this possibility further, we introduced a substitution at the dimerization interface predicted to disrupt hydrogen bonding between RefZ subunits. Substitution of an alanine at R106, an invariant residue in *Bacillus refZ* homologs that participates in two hydrogen bond contacts across the dimer interface (four bonds total)(Fig 5B), resulted in the reduced self-interaction as expected (Fig 7B).

B2H analysis of the 10 rLOF variants revealed three classes of self-interaction phenotypes: loss-of-interaction, gain-of-interaction, and wild-type interaction. Three rLOF variants, L153R, R102C, and R116W classed as loss-of-interaction. Like R106, R102 and L153 are located on the dimer interface. R102 contributes a total of two hydrogen bonds to RefZ dimer formation (Fig 5B). Substitution of a cysteine at R102 would therefore be expected to reduce dimerization and this is consistent with the reduced self-interaction observed (Fig 7B). The L153R substitution introduces a longer, positively charged side chain into a hydrophobic region of the RefZ dimer interface, and thus is also predicted to reduce dimerization (Fig 5A). No self-interaction was observed for the L153R variant, consistent with the structural prediction. These data suggest that the loss-of-function phenotypes of R102C and L153R may be related to a reduced ability to dimerize.

Three variants, E53K, R116S, and E179K displayed enhanced self-interaction compared to wild type (Fig 7B). E53K is positioned on α4, the helix connecting the regulatory domain (α4-α10) to the DNA-binding domain (α1-α3). In TetR and QacR, conformational changes caused by ligand binding to the regulatory domain are transmitted through α4 to the HTH, leading to DNA release^62^. Since the E53K mutant also shows higher affinity for non-specific DNA (Fig 6), we hypothesize that E53K facilitates a conformation that both dimerizes and binds DNA more readily. Given that the R116S and R116W variants display opposite phenotypes (enhanced and weakened self-interaction, respectively), R116 clearly has an important role in determining RefZ’s dimerization state. The E179K substitution is located just proximal to α8, a helix that participates in hydrophobic interactions between RefZ subunits (Fig 3B). The E179K substitution may cause a change in RefZ’s overall conformation that enhances hydrophobic interactions between helices α8 and α8’ of the RefZ subunits.

Four variants, R102S, E61K, E117D, and E117G, exhibited self-interaction comparable to wild type (Fig 7B). Notably, even though the R102S and E117D substitutions support wild-type self-interaction (Fig 7B) and *RBM* binding (Fig 6), they are not functional *in vivo*. These results suggest that R102 and E117 are perturbed in functions not revealed by the *ex vivo* assays. At the same time, six of the 10 rLOF variants display either reduced or increased self-interaction, suggesting that the ability of RefZ to switch between monomer and dimer forms is likely important for the mechanism leading to FtsZ inhibition.

### Thermostability of RefZ and the rLOF variants

To examine the effect of the rLOF substitutions on RefZ’s thermostability, we performed differential scanning fluorimetry (DSF). Wild-type RefZ displayed a single transition melting curve (S5 Fig, WT), with a melting temperature (Tm) of 39°C (Fig 7C). With the exception of R116W, all of the variants displayed single transition melting curves (S5 Fig). Most of the variants exhibited a lower Tm compared to wild type (L153R<R102C<R116S<R102S<WT). Notably, L153R and R102C were the most destabilized (−5°C and −4°C, respectively) and also showed the weakest self-interaction in the B2H (Fig 7B). Conversely, E53K was more thermostable than wild type and also displayed the most self-interaction by B2H (Fig 7C). R116W also displayed reduced thermostability and self-interaction; however, unlike L153R and R102C, the R116W melting curve displayed two transitions, suggesting that the R116W variant assumes more than one conformation in solution. These results suggest that RefZ and the rLOF variants may assume multiple conformations in solution, and that RefZ’s oligomerization state may be partly reflected in the thermostability measurements.

## Discussion

RefZ is required for the timely redistribution of FtsZ from midcell to the pole^49^. RefZ can also inhibit Z-ring assembly and filament cells when it is artificially induced during vegetative growth, an activity that requires DNA binding^49^. Under its native regulation, RefZ is expressed early in sporulation and requires the *RBMs* to facilitate precise capture of the chromosome in the forespore^48^. Together, these results suggest that RefZ’s effect on FtsZ, whether direct or indirect, is regulated by interactions with the nucleoid. Strikingly, the *RBMs* and their relative positions on the chromosome with respect to *oriC* are conserved across the entire *Bacillus* genus, indicating there is strong selective pressure to maintain the location of the *RBMs*. In *B. subtilis*, the *RBMs* are positioned in the cell near the site of polar septation. These observations, and the fact that RefZ, like SlmA (the NO protein of *E. coli*) belongs to the TetR family of DNA-binding proteins led us to hypothesize that RefZ binds to the *RBMs* to tune Z-ring positioning relative to the chromosome during sporulation.

To determine if RefZ’s FtsZ-inhibitory activity was important for chromosome capture, we took advantage of RefZ’s vegetative misexpression phenotype (filamentation and cell killing in a sensitized background) to isolate 10 rLOF variants capable of binding DNA, but unable to inhibit FtsZ. All 10 of the rLOF variants were unable to support correct chromosome capture (Fig 2), consistent with a model in which RefZ-RBM complexes act through FtsZ to facilitate precise septum placement with respect to the chromosome during polar division. This model is also supported by recent evidence showing that on average, Δ*refZ* mutants position Z-rings approximately 15% further away from the cell pole compared to the wildtype^67^.

### RefZ and SlmA do not inhibit FtsZ through a common mechanism

To better understand RefZ’s mechanism of action at the molecular level, wildtype RefZ and the rLOF variants were overexpressed, purified, and analyzed using structural and biochemical approaches (summarized in Table 2). The RefZ crystal structure revealed that RefZ is capable of forming a homodimer (Fig 3), similar to other TetR proteins, including SlmA. The relative locations and nature of the loss-of-function substitutions in RefZ and SlmA are different (Fig 4), suggesting that if RefZ interacts with FtsZ directly, then RefZ’s mechanism of action is distinct from that of SlmA. At least some mechanistic differences would be expected, as the C-terminal tails of FtsZ from *B. subtilis* and *E. coli* are distinct. More specifically, while the portion of *E. coli* FtsZ observed to interact with SlmA in the co-crystal is relatively conserved (DIPAFLR in *E. coli* and DIPTFLR in *B. subtilis)*, the remainder of the C-termini differ significantly (KQAD in *E. coli* and NRNKRG in *B. subtilis)*.

**Table 2.**
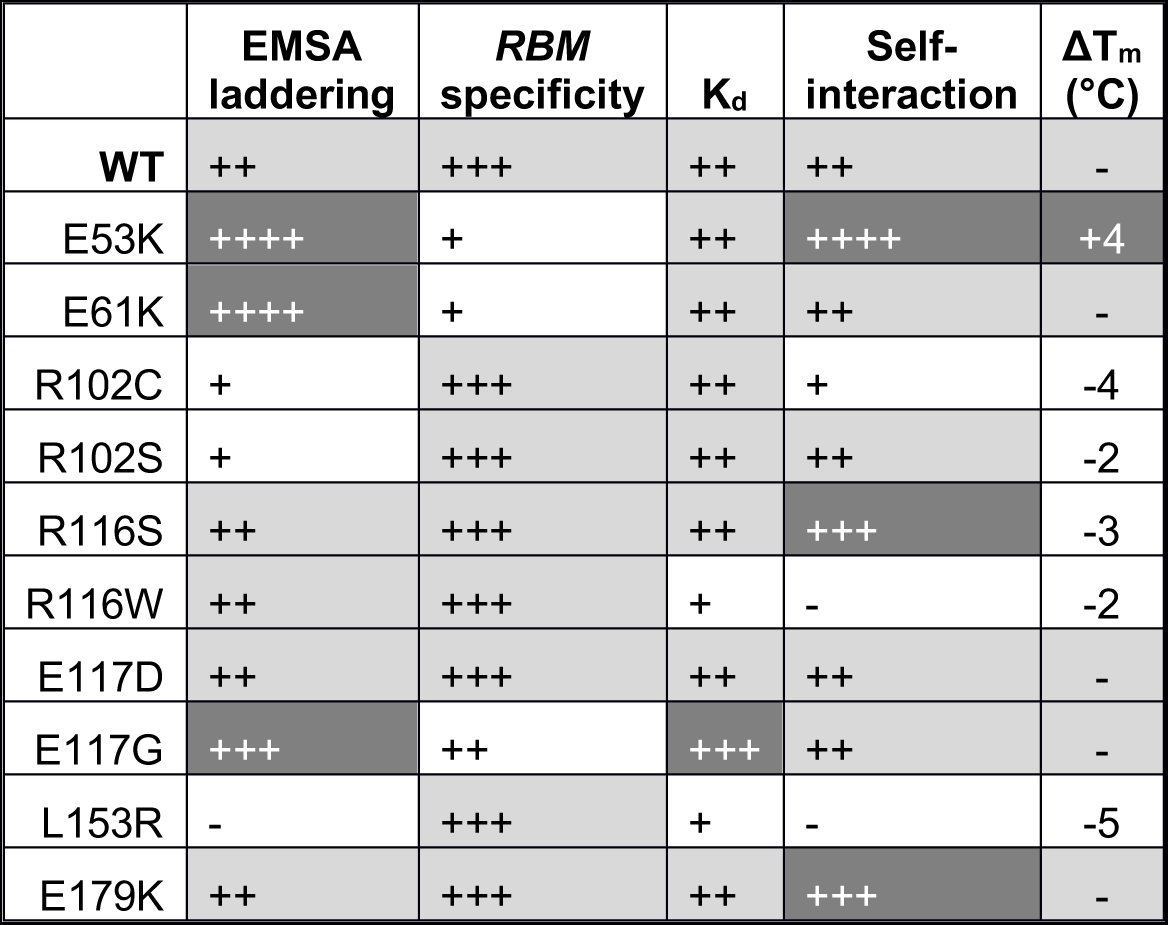
Summary of rLOF phenotypes.

|  | **EMSA**<br>**laddering** | ***RBM***<br>**specificity** | **K<sub>d</sub>** | **Self-**<br>**interaction** | **ΔT<sub>m</sub>**<br>**(°C)** |
| --- | --- | --- | --- | --- | --- |
| **WT** | ++ | +++ | ++ | ++ | - |
| E53K | ++++ | + | ++ | ++++ | +4 |
| E61K | ++++ | + | ++ | ++ | - |
| R102C | + | +++ | ++ | + | -4 |
| R102S | + | +++ | ++ | ++ | -2 |
| R116S | ++ | +++ | ++ | +++ | -3 |
| R116W | ++ | +++ | + | - | -2 |
| E117D | ++ | +++ | ++ | ++ | - |
| E117G | +++ | ++ | +++ | ++ | - |
| L153R | - | +++ | + | - | -5 |
| E179K | ++ | +++ | ++ | +++ | - |

### The role of self-interaction and RBM-binding in RefZ function

An important finding of this study is that both enhanced and reduced RefZ dimerization are correlated with loss-of-function phenotypes *in vivo*. B2H analysis indicates that the majority of rLOF variants (6/10) exhibited either stronger or weaker self-interaction (Fig 7B), suggesting that RefZ’s propensity to switch between a monomer and dimer states is integral to affecting FtsZ function. Two rLOF variants (R102C and L153R) possess substitutions predicted to disrupt dimerization (Fig 5), a result corroborated by B2H analysis (Fig 7B). L153R also causes a 2-fold reduction in affinity for *RBM*-containing DNA, which could affect its ability to appropriately localize to *RBMs in vivo*.

Two rLOF variants (E53K and E61K) are located on α4. Based on the observation that E53K and E61K exhibit enhanced laddering and an increased apparent affinity for nonspecific DNA by EMSA (Fig 6), we propose that these variants assume a conformation that is more favorable for nonspecific DNA-binding than that assumed by wild type. *In vivo*, enhanced nonspecific binding would reduce the formation of RefZ-RBM complexes, which prior data suggest is the functional form of RefZ^48,49^.

The ability of RefZ to generate DNA laddering in EMSAs (Fig 6 and S3 Fig) is presumably due to the association of additional RefZ subunits to adjacent DNA after the initial pair of dimers binds the RBM^48^. Other TetR proteins, including SlmA, have also been observed to “spread” on DNA *in vitro^58,63,68^*. In the case of SlmA, spreading on DNA is hypothesized to facilitate interaction with the exposed C-terminal tails of FtsZ to promote filament breakage^58^. Although genetic and cell biological data suggest RefZ and FtsZ interact^48,49,67^, evidence for direct interaction between RefZ and FtsZ is lacking. Attempts to test for RefZ-FtsZ interaction *in vitro* have been impeded by RefZ’s limited solubility outside of the specific conditions identified in this study. Therefore, the precise mechanism by which RefZ affects FtsZ remains to be determined.

One of the most interesting observations obtained from characterizing the rLOF variants is that the R116S and R116W substitutions on the first turn of α7 result in opposite self-interaction phenotypes (Fig 7B). Both variants behave comparably with regard to affinity and specificity for the *RBM*-containing DNA (Fig 6), suggesting the loss-of-function phenotypes are not attributable to differences in DNA interaction or protein misfolding. Instead, these results suggest that R116 is a key residue in determining the stability of the RefZ dimer. We hypothesize that R116 participates in intramolecular bonds with residues within a flexible loop region (between α6 and α7, residues 109-114)(Fig 3A), possibly contributing to the formation of a more stable homodimer. R116 could participate in formation of either ionic or hydrogen bonds with a invariant aspartate residue (D111) located in the flexible loop. Our ability to assess R116’s role in intramolecular bond formation is limited in the current crystal structure, as the electron density for the R116 side-chain is not well defined. Moreover, the electron density for the main chain of the flexible loop is moderately disordered, showing peaks of positive Fo-Fc electron density next to the I110 and D111 side-chains.

R116 is also immediately adjacent to E117, another critical residue identified in this study. E117D is the only rLOF variant that is loss of function with regard to inhibiting cell division and capturing the forespore chromosome, yet is not detectably altered in the other RefZ properties implicated in function (Table 2). If RefZ targets FtsZ directly, then these data point toward E117 as a likely candidate residue for mediating interaction. The E117D substitution is intriguing because the glutamate to aspartate change is highly conservative; however, if the interaction is direct, the shorter sidechain of the aspartate could compromise RefZ’s ability to target FtsZ. It has not escaped our attention that many regulators of FtsZ including FtsA, ZapD, and MinD possess glutamate or aspartate residues near the implicated FtsZ C-terminal tail binding site which are proceeded by either a hydrophobic or polar uncharged residue followed by an arginine or lysine. For example, FtsA from *Thermotoga maritima* possesses LRE^69^, ZapD from *E. coli* and a variety of Gammaproteobacteria I(R/K)E^70,71^, and MinD a highly conserved ARD^72^. Whether these residues represent a *bona fide* motif involved in FtsZ regulation remains to be determined, but it is intriguing that two residues identified as critical for RefZ function fall within an IRE sequence.

### Working model for RefZ-mediated septum positioning

Based on the data available, we propose a model in which RefZ mediates chromosome capture by fine-tuning the position of FtsZ assembly over the forespore-destined chromosome. In our model, RefZ is primed to inhibit FtsZ polymerization near the pole by binding specifically to the polarly-localized *RBMs*. Based on structural studies of other TetR family proteins and the observation that RefZ binds to *RBMs* in units of two and four *in vitro*^48,49^, RefZ likely binds each *RBM* as a pair of dimers. We were not able to report RefZ copy number as native RefZ levels are too close to the detection limit of our antibodies; however, our preliminary data suggest that RefZ is likely a relatively low copy number protein.

Current data suggest the activity of RefZ inhibits rather than promotes FtsZ assembly^48,49,67^. This raises the question as to how an inhibitor of FtsZ could act near the pole to promote precise placement of a polar division apparatus. In our model, RefZ is a locally-acting inhibitor of FtsZ and its primary function is not to inhibit the formation of polar Z-rings altogether, but rather to tune the location of Z-ring assembly away from the immediate vicinity of the *RBMs*. Based on comparative analysis of the rLOF mutants, both decreased and increased ability to dimerize appears to be detrimental to the inhibitory function of RefZ. This implies that a dynamic process of monomer-dimer exchange, not maintaining a specific oligomeric state, is what is important for RefZ function. One possibility is that *RBM*-bound dimers disassociate from DNA as monomers after engaging with FtsZ.

We present no evidence that RefZ’s DNA association or monomer-dimer exchange is influenced by a ligand, and no obvious ligand binding pocket is observed in the regulatory domain of the solved crystal structure. At the same time, we do not exclude the possibility that RefZ activity could be regulated through interaction with FtsZ or ligand binding. Recently EthR, an important TetR family protein from *Mycobacterium tuberculosis* that regulates drug resistance, was shown to bind the nucleotide cyclic-di-GMP^73^. Interestingly, EthR’s proposed nucleotide binding region (based on mutagenesis and docking studies) is at the dimer interface, outside the canonical ligand binding pocket^73^ (near R102 in RefZ).

Another paradox raised is why a Δ*refZ* mutant exhibits a slight delay in shifting Z-rings from midcell to the pole during sporulation^49^. If RefZ acts as an inhibitor at the pole, then assembly of the polar Z-ring would be expected to accelerate in a Δ*refZ* mutant. This seeming contradiction may be explained by considering RefZ’s localization during sporulation. At early timepoints, just before polar division occurs, RefZ-GFP localizes as foci near the poles. These foci likely represent RefZ-RBM complexes, as they are lost in a RefZ mutant that cannot bind DNA^49^. Around the time polar division initiates, the polar RefZ foci become less apparent and RefZ is observed to coalesce near midcell at or near the membrane^49^. The redistribution of RefZ’s inhibitory activity from the pole to midcell as sporulation progresses could facilitate disassembly of the midcell Z-ring and its reassembly at the pole^42,43^. Preliminary data also suggest that RefZ has a second role, to prevent additional midcell divisions as sporulation progresses (Miller and Herman, unpublished), and current investigations are aimed at exploring this possibility.

## Methods

### General methods

Strains, plasmids, and oligonucleotides are listed in Supplemental S1, S2, and S3 Tables, respectively. All *Bacillus subtilis* strains were derived from *B. subtilis* 168 or PY79. Strain and plasmid construction is detailed in the Supporting Information. Transformations in *B. subtilis* were carried out using a standard protocol as previously described ^74^ unless otherwise stated. For selection in *B. subtilis*, antibiotics were included at the following concentrations: 100 μg ml^−1^ spectinomycin, 7.5 μg ml^−1^ chloramphenicol, 10 μg ml^−1^ kanamycin, 10 μg ml^−1^ tetracycline, 0.8 μg ml^−1^ phleomycin, and 1 μg ml^−1^ erythromycin (erm) plus 25 μg ml^−1^ lincomycin (MLS). For transformation and selection in *E. coli*, antibiotics were included at the following concentrations: 100 μg ml^−1^ ampicillin, 25 μg ml^−1^ kanamycin, and 25 μg ml^−1^ chloramphenicol (for protein overexpression). Co-transformations for B2H assays were selected for on LB plates supplemented with 50 μg ml^−1^ ampicillin, 25 μg ml^−1^ kanamycin, and 0.2% (v/v) glucose.

### Two-step genetic selection-screen to isolate rLOF mutants

Comprehensive details on construction of the Gibson assemblies and strains below are available in the supplemental text. The *refZ* gene was mutagenized by error-prone PCR and the mutant fragment library was introduced into an IPTG-inducible misexpression construct using Gibson assembly^53^. Multiple assembly reactions were pooled on ice and directly transformed into super-competent BAM168 cells (selection-screen background). For transformations, competent cell aliquots were thawed at room temperature and 0.2 ml were incubated in a 13 mm glass test tube with 20 μl assembly reactions for 90 min in a rollerdrum at 37°C before selecting on LB plates supplemented 100 μg ml^−1^ spectinomycin and 1 mM IPTG. After overnight growth at 37°C, surviving transformants were patched on LB plates supplemented with 1% (w/v) starch to screen for integration at *amyE*, and on LB plates supplemented with the following antibiotics to assess the presence of the expected parental background resistances: 7.5 μg ml-1 chloramphenicol, 10 μg ml-1 kanamycin, 10 μg ml-1 tetracycline, and 1 μg ml-1 erythromycin (erm) plus 25 μg ml-1 lincomycin (MLS). Transformants were also patched on LB plates supplemented with 100 μg ml^−1^ spectinomycin and 1 mM IPTG and 40 μg ml^−1^ X-gal to screen for *lacZ* expression from the P_*spremo*_ promoter. Replica plates were grown overnight at 37°C. Surviving *rLOF* mutants that did not turn blue on patch plates were cultured from replica plate in liquid LB and stored at −80°C. Genomic DNA prepared from these strains was PCR amplified with OJH001 and OJH002 to test for the presence of the expected integration product. PCR products of the expected size were sequenced to identify mutations.

### Generation of super-competent cells

Super-competency was achieved using two-fold approach to maximize transformation efficiency. First, BAM168 (selection-screen background) harbors a xylose-inducible copy of *comK* at the non-essential *lacA* locus^75^. The presence of 1% (w/v) xylose in standard transformation cultures improved efficiency ~2.5-fold compared to cultures grown without xylose. Second, competent cells were prepared by modifying an established^74^ two-step *B. subtilis* competent cell protocol as described below. The modifications improved transformation efficiency an additional 7-fold over xylose induction alone. A single colony of freshly streaked recipient cells (BAM168) was used to inoculate a 250 ml baffled flask containing 25 ml of 1X MC medium (10.7 g L^−1^ K_2_HPO_4_, 5.2 g L^−1^ KH_2_PO_4_, 20 g L^−1^ glucose, 0.88 g L^−1^ tri-sodium citrate dihydrate, 0.022 g L^−1^ ferric ammonium citrate, 1 g L^−1^ casein hydrolysate (Neogen), 2.2 g L^−1^ potassium glutamate monohydrate, 3 mM MgSO4, and 0.02 g L^−1^ L-Tryptophan)^74^. The culture was grown overnight (20-22 h) in a 37°C shaking waterbath set at 250 rpm. The overnight culture (OD_600_ 1.5-2.5) was diluted to an OD_600_ of 0.1 in a 250 ml baffled flask containing 40 ml of 1X MC supplemented with 1% (w/v) xylose. The culture was incubated at 37°C in a shaking waterbath set at 200 rpm. After 5-6 h of growth, the OD_600_ was monitored every 30 min until readings remained unchanged between two timepoints, at which point the culture was diluted 1:10 with pre-warmed 1X MC supplemented with 1% (w/v) xylose to a final volume of 250 ml in a 2 L flask. After 90 min of growth at 37°C and 280 rpm, cells were harvested at room temperature at 1,260 x *g* for 10 min in six 50 ml conical tubes. Twenty ml of the culture supernatant was retained and mixed with 5 ml 50% (v/v) glycerol. The diluted supernatant was used to gently resuspend the pellets, and the cell suspensions were immediately frozen at - 80°C in aliquots.

### Blue-white screen to assess RBM-binding by rLOF mutants

Misexpression constructs harboring either wild-type *refZ* (BAM374), *rLOF* mutants (BAM400, 403, 407, 409, 411, 440, 443, 444, 449, 462), or an empty Phy vector (BAM390) in clean selection-screen backgrounds (Supplemental Text) were streaked from frozen glycerol stocks on LB plates supplemented with 100 μg ml^−1^ spectinomycin and 0.2% (v/v) glucose and grown overnight at 37°C. Single colonies were used to inoculate 3 ml of Lysogeny Broth (LB-Lennox) and cultures were grown in a rollerdrum at 30°C until early to mid-log (3-5 h). Cultures were normalized to the lowest OD_600_ with PBS (10^0^) and serially diluted (10^−1^, 10^−2^,10^−3^). Five μl of each dilution was spotted on LB plates supplemented with 100 μg ml^−1^ spectinomycin and 1 mM IPTG and 40 μg ml^−1^ X-gal followed by overnight incubation at 37°C to visually screen for *lacZ* expression from the P_*spremo*_ promoter. Plates were scanned with a ScanJet G4050 flatbed scanner (Hewlett Packard) using VueScan software and medium format mode. Images were processed using Adobe Photoshop (version 12.0).

### Misexpression of wild-type refZ and rLOF variants

Misexpression constructs harboring either wild-type *refZ* (BJH228) or the *rLOF* mutants (BAM428, 431, 434, 436, 450, 451, 454, 455, 457, 490) in a wild-type background (Supplemental Text) were streaked from frozen glycerol stocks on 100 μg ml^−1^ spectinomycin plates and grown overnight at 37°C. CH cultures (25 ml) were prepared as described under *Fluorescence microscopy*. Misexpression was induced with 1 mM IPTG following 1.5-2 h of growth at 37°C (approx. OD_600_ 0.10). For the uninduced controls in Figure 1C and 1D, an independent culture of the control strain, BJH228 (P_*hy*_-*refZ*), was grown in parallel but was not induced. Growth was resumed at 37°C with shaking for 45 min (see *Western blotting*) or 90 min (see *Fluorescence microscopy*) before 1 ml samples were harvested.

### Fluorescence microscopy

For microscopy experiments, isolated colonies were used to inoculate 5 ml CH and cultures were grown overnight at room temperature in a rollerdrum. Cultures below an OD_600_ of 0.7 were used to inoculate 25 ml CH medium in 250 ml baffled flasks to a calculated OD_600_ of 0.006 (for misexpression) or 0.018 (for chromosome capture assays) and cultures were grown for the indicated time at 37°C in a shaking waterbath set at 280 rpm. Samples were collected at 6,010 x *g* for 1 min in a tabletop microcentrifuge. Following aspiration of supernatants, pellets were resuspended in 3-5 μL of 1X PBS containing 0.02 mM 1-(4-(trimethylamino)phenyl)-6-phenylhexa-1,3,5-triene (TMA-DPH)(Life Technologies) and cells were mounted on glass slides with polylysine-treated coverslips. Images were captured and analyzed with NIS Elements Advanced Research (version 4.10) software, using 600 ms (CFP), 900 ms (YFP), or 1 s (TMA) exposure times on a Nikon Ti-E microscope equipped with a CFI Plan Apo lambda DM 100X objective, a Prior Scientific Lumen 200 Illumination system, C-FL UV-2E/C DAPI, C-FL YFP HC HISN Zero Shift, and C-FL Cyan GFP filter cubes, and a CoolSNAP HQ2 monochrome camera.

### Western blotting

Samples were harvested at 21,130x *g* for 1 min in a tabletop centrifuge. Pellets were washed with 50 μl of 1X PBS and the remaining supernatant was carefully removed using a P20 pipet. Pellets were frozen at −80°C until processing. Frozen pellets were thawed on ice before resuspension in 25 μl of lysis buffer (20 mM Tris [pH 7.5], 10 mM EDTA, 1 mg ml^−1^ lysozyme, 10 μg ml^−1^ DNase I, 100 μg ml^−1^ RNase A, and 1 mM phenylmethylsulfonyl fluoride). Samples were normalized by OD_600_ values obtained at the time of harvest by diluting resuspensions in additional lysis buffer before incubating at 37°C for 15 min. Samples were diluted 1:1 with 2X sample buffer (250 mM Tris [pH 6.8], 10 mM EDTA, 4% (v/v) SDS, 20% (v/v) glycerol, and 10% (v/v) 2-mercaptoethanol) and boiled for 10 min. Five μl of each lysate was loaded on a 4-20% gradient polyacrylamide gel (Lonza) and proteins were separated by electrophoresis prior to transfer to a nitrocellulose membrane (Pall)(1 h at 60 V). Membranes were blocked for 1 h at room temperature in 5% (w/v) nonfat milk in PBS [pH 7.4] with 0.05% (v/v) Tween-20. Membranes were incubated overnight at 4°C with polyclonal rabbit anti-RefZ antibody (Covance) diluted 1:1,000 in 5% (w/v) nonfat milk in PBS [pH 7.4] with 0.05% (v/v) Tween-20. Membranes were washed prior to a 1 h room temperature incubation with horseradish peroxidase-conjugated goat anti-rabbit Immunoglobulin G secondary antibody (Bio-Rad) diluted 1:10,000 in 5% (w/v) nonfat milk in PBS [pH 7.4] with 0.05% (v/v) Tween-20. Washed membranes were incubated with SuperSignal West Femto Maximum Sensitivity substrate (Thermo Scientific) according to the manufacturer’s instructions. Chemiluminescence was detected and imaged using an Amersham Imager 600 (GE Healthcare). Images were processed using ImageJ64^76^.

### Chromosome capture assay with the rLOF mutants

Strains used in the chromosome capture assay in Fig 2 harboring the left arm (61° P_*spoIIQ*_-*cfp*) or right arm (+51° P_*spoIIQ*_-*cfp*) reporter in the wild type, *refZ* mutant, or *rLOF* mutant trapping backgrounds (Supporting Information S1 Table) were streaked from frozen stocks on LB agar plates and grown overnight at 37°C. Chromosome capture assays were carried out as previously described^18,48^. CH cultures (25 ml) were prepared as described in *Fluorescence microscopy* and grown for 2.5-3 h (OD_600_ 0.6-0.8) before sporulation was induced by resuspension according to the Sterlini-Mandelstam method^74^. Growth was resumed at 37°C in a shaking waterbath for 2.5 h prior to TMA-DPH, YFP, and CFP image acquisition (see *Fluorescence microscopy)*.

Each strain harbors a σ^F^-dependent oriC-proximal reporter (−7° P_*spoIIQ*-*yfp*_) that is captured in the forespore in 99.5% of sporulating cells. Cells expressing YFP serve as the baseline for total sporulating cells counted in the field. To visualize cells in a given field that expressed the left or right arm reporters in the forespore, captured YFP and CFP images were individually merged with the TMA (membrane) image. The total number of forespores with YFP signal (total YFP) or CFP signal (total CFP) were manually marked and counted as described previously^48^.

For quantitation and statistical analysis, a minimum of 1,500 cells per strain were counted from three independent biological and experimental replicates, with the exception of wildtype (left and right arms, n=7) and the E53K (right arm, n=4). The average proportion of cells expressing both reporters for each strain is given in Figure 2, with error bars representing one standard deviation above and below the average. Twotailed Student’s t-tests were performed to determine the P-values indicated in the pairwise comparisons.

### Protein Purification

*E. coli* BL21(DE3) pLysS competent cells were transformed with either pLM025a (RefZ-His6) or pEB013-pEB022 (rLOF-His6) and grown overnight at 37°C on LB plates supplemented with 25 μg ml^−1^ kanamycin, 25 μg ml^−1^ chloramphenicol and 0.1% (v/v) glucose. Transformants were scraped from plates and resuspended in 2 ml of Cinnabar High-Yield protein expression media (Teknova, Cat No. 3C8488) containing 25 μg ml^−1^ kanamycin, 25 μg ml^−1^ chloramphenicol and 0.1% (v/v) glucose. The OD_600_ was measured and used to inoculate 4 x 25 ml of the same medium in 250 ml baffled flasks to an OD_600_ of 0.1. Cultures were grown at 37°C in a shaking waterbath at 280 rpm for 6-7 h until the culture density reached OD_600_ = 5.0. Protein expression was induced with 1 mM IPTG and growth was resumed for an additional 3 h before cultures were harvested by centrifugation at 9,639 x *g* for 5 min at 4°C. Pellets were stored at −80°C until processing. Four pellets (25 ml culture each) were resuspended in 40 ml of lysis Buffer (50 mM Tris-HCl [pH 9.0], 300 mM KCl, 10% (v/v) glycerol, and 10 mM imidazole). 1 μl protease inhibitor (Sigma-Aldrich, Cat No. P8465)(215 mg powder dissolved in 1 ml of DMSO and 4 ml ddH_2_0) was added per 35 OD_600_ units. DNase I was added to a final concentration of 1 μg ml^−1^ of cell suspension. Suspensions were passed through a Microfluidizer LM20-30 five times at 10,000 psi. Cell debris was cleared by centrifugation at 22,662 x *g* for 30 min at 4°C. Supernatants were passed over a 1 ml bed volume of Nickel-NTA agarose beads (Qiagen, Cat No. 30210) preequilibrated with lysis buffer. Bound protein was washed with 10 ml of wash buffer (50 mM Tris-HCl [pH 9.0], 300 mM KCl, 10% (v/v) glycerol, and 20 mM imidazole). Protein was eluted with 7 ml of elution buffer (50 mM Tris-HCl [pH 9.0], 300 mM KCl, 10% (v/v) glycerol, and 250 mM imidazole) and collected as ~250 μl fractions. 2 μl was removed from each fraction for SDS-PAGE analysis, and elutions were immediately stored at - 80°C. Peak elution fractions were thawed and pooled before dialyzing at 4°C with stirring into either elution buffer (50 mM Tris-HCl [pH 9.0], 300 mM KCl, 10% (v/v) glycerol, and 250 mM imidazole) or ddH2O using Slide-A-Lyzer^®^ 7.0 kDa MWCO dialysis cassettes (Thermofisher) Scientific). Final protein concentrations were determined using Bradford reagent (Bio-Rad) and a BSA standard.

### Protein crystallization, data collection, and data analysis

RefZ-His6 was overexpressed and purified as described above. Before dialysis the RefZ concentration was determined and dsDNA (generated by annealing OEB025/OEB026) was added to a 4:1 molar ratio of *RefZ:*RBM_L2_*-24bp*. The protein was dialyzed into 50 mM Tris-HCl [pH 8.5] and 300 mM KCl. After dialysis, RefZ was concentrated in a 10 kDa Vivaspin Turbo MWCO filter (Sartorius) to ~5 mg ml^−1^, and 0.5-1.0 μl of the concentrated protein was used to set crystallization plates. RefZ crystals formed within 48 h by hanging drop vapor diffusion at 16°C after mixing the protein in a 1:1 volume ratio with 10% ethanol (v/v), 0.1 M imidazole [pH 8.0], and 0.2 M MgCl_2_. The crystals were cryoprotected in 20% (v/v) glycerol in mother liquor before flash freezing in liquid nitrogen. For anomalous signal, RefZ crystals were soaked with 1 mM lead acetate for 5 h and the data were collected at the Argonne National Lab APS synchrotron, beamlines 23-ID, at 0.9496 Å. Diffraction data were indexed, integrated, and scaled in HKL2000^77^ and the single heavy atom site was identified by phasing using single anomalous dispersion (SAD) in the SHELX program^78^. The resultant phases were extended to a native crystal data set collected at the same beamline at 0.98 Å. The native set was indexed, integrated, and scaled using PROTEUM3 software (Version 2016.2, Bruker AXS Inc). The native crystal data were truncated in Ctruncate^79^ from CCP4 suite^80^ and subjected to iterative building and phase improvement by PHENIX^81^. The partial model produced by PHENIX was rebuilt in BUCCANEER^82^ relying on improved phases. BUCCANEER was able to build the whole model in one continuous chain, docked in sequence and covering residues 1-200. The model was improved through iterative runs of inspection and manual modification in COOT^83^ and refinement in PHENIX^81^ with simulated annealing on initial runs. The data collection and refinement statistics can be found in Table 1.

### Annealing of oligos to generate dsDNA

Oligonucleotides were resuspended in annealing buffer (10 mM Tris-HCl [pH 7.5], 50 mM NaCl, and 1 mM EDTA) to a concentration of 1 mM. Equal volumes were mixed and annealed in a thermocycler by heating to 95°C for 2 min followed by ramp cooling for 45 min to 25°C. The annealing buffer was removed by dialysis into ddH2O with Slide-A-Lyzer^®^ 7.0 kDa MWCO Dialysis Cassettes (Thermo Scientific).

### Electrophoretic gel mobility shift assays

DNA fragments centered on either the native (using *B. subtilis* 168 as template) or the mutant (using BJH205 as template) *RBM_L1_* sequence^48^ were generated by PCR using primer pair OEB009 and OEB010. Purified RefZ-His6 or rLOF-His6 protein (final concentrations indicated in Figure 6) were incubated with 10 nM *RBMl1* or *RBML1mu* DNA probes in binding buffer (150 mM KCl and 10 mM Tris-HCl [pH 8.0]) for 30 min. After 30 min incubation, 10X loading buffer (50 mM EDTA [pH 8.0], 1 mM Tris-HCl [pH 8.0] and 45% (v/v) glycerol) was added to a final concentration of 1X and binding reactions were resolved at room temperature on a 5% TBE polyacrylamide gel run for 45 min at 150 V (Fig 6) or a 7.5% TBE polyacrylamide gel for 17 min at 200 V (S3 Fig). After electrophoresis, gels were incubated with agitation in 1X SYBR Green EMSA gel stain (Life Technologies)(diluted from 10,000X stock in TBE buffer) for 5 min then rinsed with dH2O. Stained DNA was imaged with a Typhoon FLA 9500 scanner using the setting for Fluorescence and LPB (510LP) filter for SYBR Green. The data presented in Figure 6 is representative of a minimum of three independent experimental replicates for wild type and each variant.

### Bio-layer Interferometry Assay

The Octet system (Pall Forte Bio) was used to monitor the kinetic interactions between wild-type RefZ or the rLOF variants and *RBM*-containing DNA. Streptavidin biosensors (Part NO 18-5019) were purchased from Pall Forte Bio. A 41 bp *RBM*-containing (*RBM_L1_*) segment of dsDNA was generated by annealing 5’ biotinylated OEB091 with OEB092 as described (see *Annealing of oligos to generate dsDNA*) except that the annealing buffer was not removed by dialysis. All subsequent assays were performed in DNA binding buffer (150 mM KCl and 10 mM Tris-HCl [pH 8.0]). Sensors were pre-equilibrated for 10 min at room temperature in DNA-binding buffer to establish a baseline reading. Sensors were then dipped into a well containing 50 nM *RBMli* dsDNA and incubated for 2 min with shaking at 1,000 rpm to immobilize DNA on the biosensor. The sensor was washed for 30 sec to establish a new baseline before transfer to a solution containing 800 nM of wild-type RefZ or rLOF variants. Following a 3 min monitored association, the complex was placed into fresh buffer and dissociation was monitored continuously for 15 min. The K_d_ was calculated using the global fit in Pall Forte Bio’s analysis software. Three experimental replicates of each assay were performed except for variant R102C (n=4). The mean values and standard deviations are given in Figure 6. P-values were determined using a two-tailed unpaired Student’s t-test.

### Size-exclusion chromatography

A Superdex 200 PC 3.2/30 3.2 × 300 mm column was equilibrated with 50 mM Tris-HCl [pH 9.0], 300 mM KCl, and 10% (v/v) Glycerol. Wild-type RefZ and rLOF proteins from frozen stocks (ddH_2_O) were diluted to a final concentration of 1 mg ml^−1^ in 200 μl of buffer (50 mM Tris-HCl [pH 9.0], 300 mM KCl, 10% (v/v) Glycerol). Samples were pre-spun at 21,130 x *g* for 10 min at 4°C in a tabletop centrifuge prior to injection. The absorbance at 280 nm was continuously measured and the V_e_, peak maximum, was taken from the resulting elution profile and used to calculate K_av_ using the formula (V_e_ - V_o_)/(V_t_ - V_o_). The void volume, V_o_ was experimentally determined to be 7 ml. The total volume, V_t_, of the column was 24 ml. The apparent molecular mass was estimated using a curve generated from an identical run with a molecular mass standard (Bio-Rad Gel filtration chromatography standard, cat. no. 151-1901).

### Bacterial 2-hybrid analysis of rLOF variants

Assays were carried out essentially as previously described^48,66^. Plasmids harboring wild-type *refZ* and the rLOF sequences fused with C-terminal T18 and T25 tags (see Supplemental for plasmid construction) were co-transformed into competent *E.coli* DHP1 (_cya_-) cells with selection on LB plates supplemented with 50 μg ml^−1^ ampicillin, 25 μg ml^−1^ kanamycin, and 0.2% (v/v) glucose. Co-transformed *E.coli* strains were streaked from frozen stocks and single colonies were cultured in 4 ml of LB supplemented with 50 μg ml^−1^ ampicillin, 25 μg ml^−1^ kanamycin, and 0.1% (v/v) glucose in a 37°C roller drum to mid-log growth phase. Culture samples were normalized to the lowest OD culture with fresh LB supplemented with 50 μg ml^−1^ ampicillin and 25 μg ml^−1^ kanamycin, and 5 μl were spotted on M9-glucose minimal plates supplemented with 50 μg ml^−1^ ampicillin, 25 μg ml^−1^ kanamycin, 250 μM IPTG, and 40 μg ml^−1^ X-gal. Pairwise interactions between the T18 and T25 fusions were assessed by monitoring the development of blue color (corresponding to *lacZ* expression) following 40-50 h of growth at room temperature. Figure 7B is representative of three independent biological and experimental replicates.

### Differential Scanning Fluorimetry (DSF)

Purified RefZ or rLOF variants from frozen stocks (50 mM Tris-HCl [pH 9.0], 300 mM KCl, 10% (v/v) glycerol, and 250 mM imidazole) were thawed and diluted in 20 mM Tris-HCl [pH 7.5] to a final concentration of 10 μM. To ensure an identical final concentration of storage buffer for all rLOF variants, reactions were normalized to the maximum required concentration of storage buffer determined by the lowest rLOF variant concentration; the final buffer concentration was 0.16X. All reactions contained 5X SYPRO™ Orange Protein Gel Stain (Thermofisher) diluted to a working concentration in DMSO. The DSF assays were performed in a 96-well hardshell PCR plate (Bio-Rad, HSP9601) using a CFX96 Touch™ Real-Time PCR Detection System (Bio-Rad). The reactions were ramped from 25°C to 95°C at a rate of 1°C min^−1^.

## Accession codes

The coordinates and structure factors for RefZ have been deposited in the Protein Data Bank (PDB: 6MJ1).

## Acknowledgements

We thank Larry Dangott and the Protein Chemistry Lab at Texas A&M for helpful advice regarding protein purification, Ann Tran for her efforts toward constructing B2H plasmids and quantifying rLOF trapping data, and members of the Herman Lab for critical reading of the manuscript. This work was supported by a grant from the National Science Foundation to J.K.H. (MCB-1514629).

## Supplementary Information

**S1 Figure. Superimposition of the N-terminal domains of RefZ and QacR.** (A) Superimposition of the HTH domains of RefZ (cyan) and QacR (orange)(PDB: 1JT6)^4^. The Y43 residue on α3 of RefZ, which is required for DNA binding and the corresponding residue in QacR (Y41) are shown as sticks. (B) Superimposition of RefZ dimer (cyan) with the QacR dimer (orange) bound to *IR1* DNa (white)(PDB: 1JT0)^5^. (C) Superimposition of the HTH domains of RefZ (cyan) with QacR (orange) bound to *IR1* DNA (white)(PDB 1JT0).

**S2 Figure. Example purification profiles of wild-type RefZ and rLOF variants.** The top gel was loaded with 5 μg protein/lane and stained with coomassie blue dye (R-250). Gels below show example elution profiles from Nickel-NTA agarose beads. The elution gels were stained with coomassie brilliant blue dye (colloidal coomassie, G-250). G-250 is approximately 10 times more sensitive than R-250, allowing for detection of less abundant proteins.

**S3 Figure. EMSA laddering behavior of wild-type RefZ and rLOF variants.** (A) Laddering of DNA in the EMSAs can be observed for wild-type RefZ and to a greater extent E53K when samples are resolved at 200 V on a 7.5% TBE gel. (B)The rLOF variants R102C, R102S, and L153R do not exhibit laddering when samples are resolved at 200 V on a 7.5% TBE gel. (C) Typical bio-layer interferometry binding curve for wildtype RefZ with *RBM*-containing DNA. Sensors are pre-equilibrated for 10 min in DNA binding buffer (150 mM KCl and 10 mM Tris [pH 8]) at room temperature (not shown). The experiment is then initiated and performed at 30°C to establish a 30 sec baseline. The streptavidin sensor is dipped into a solution of biotinylated dsDNA (a 41 bp segment centered on *RBMli*) for 2 min. After incubation a new baseline is established by returning the biosensor to the DNA-binding buffer for 30 sec. The biosensor is then moved to a well containing 800 nM protein for 3 min to monitor association. The sensor is then transferred to a well containing fresh DNA-binding buffer to monitor dissociation for 15 min.

**S4 Figure. Size-exclusion chromatogram for WT RefZ.** An example Superdex 200 elution profile for 200 μl of 1 μg ml^−1^ RefZ-His6 (7.7 nmol) ran with 50 mM Tris-HCl [pH 9], 300 mM KCl and 10% (v/v) glycerol. Absorbance at 280 nm is shown on the Y-axis (mAU - milliabsorbance units). Aggregated RefZ elutes at 7.6 ml, near the column void volume (7.0 ml).

**S5 Figure. Thermostability of RefZ and the rLOF variants.** DSF estimates of wildtype RefZ-His6 and rLOF-His6 variant stability reported by fluorescence of SYPRO orange as a function of increasing temperature. (A) Representative sigmoidal melting curves. (B) Tm values (inset) were calculated by determining the temperature at which the first derivative of the fluorescence is at a minimum. ΔTm (inset) is the difference between the wild-type RefZ and each rLOF variant. Differences less than 1.5°C were not considered to be significant and are shown as dashes.

## References

1. Antonny, B. Mechanisms of membrane curvature sensing. Annu Rev Biochem 80, 101–23 (2011).

2. Updegrove, T.B. & Ramamurthi, K.S. Geometric protein localization cues in bacterial cells. Curr Opin Microbiol 36, 7–13 (2017).

3. Bramkamp, M. et al. A novel component of the division-site selection system of *Bacillus subtilis* and a new mode of action for the division inhibitor MinCD. Mol Microbiol 70, 1556–69 (2008).

4. Eswaramoorthy, P. et al. Cellular architecture mediates DivIVA ultrastructure and regulates Min activity in *Bacillus subtilis*. MBio 2(2011).

5. Gregory, J.A., Becker, E.C. & Pogliano, K. *Bacillus subtilis* MinC destabilizes FtsZ-rings at new cell poles and contributes to the timing of cell division. Genes Dev 22, 3475–88 (2008).

6. Marston, A.L. & Errington, J. Selection of the midcell division site in *Bacillus subtilis* through MinD-dependent polar localization and activation of MinC. Mol Microbiol 33, 84–96 (1999).

7. Patrick, J.E. & Kearns, D.B. MinJ (YvjD) is a topological determinant of cell division in *Bacillus subtilis*. Mol Microbiol 70, 1166–79 (2008).

8. Du, S. & Lutkenhaus, J. Assembly and activation of the *Escherichia coli* divisome. Mol Microbiol 105, 177–187 (2017).

9. Surovtsev, I.V. & Jacobs-Wagner, C. Subcellular organization: a critical feature of bacterial cell replication. Cell 172, 1271–1293 (2018).

10. Bowman, G.R. et al. A polymeric protein anchors the chromosomal origin/ParB complex at a bacterial cell pole. Cell 134, 945–55 (2008).

11. Ebersbach, G., Briegel, A., Jensen, G.J. & Jacobs-Wagner, C. A self-associating protein critical for chromosome attachment, division, and polar organization in *Caulobacter*. Cell 134, 956–68 (2008).

12. Lim, H.C. et al. Evidence for a DNA-relay mechanism in ParABS-mediated chromosome segregation. Elife 3, e02758 (2014).

13. Surovtsev, I.V., Lim, H.C. & Jacobs-Wagner, C. The slow mobility of the ParA partitioning protein underlies its steady-state patterning in *Caulobacter*. Biophys J 110, 2790–2799 (2016).

14. Badrinarayanan, A., Le, T.B. & Laub, M.T. Bacterial chromosome organization and segregation. Annu Rev Cell Dev Biol 31, 171–99 (2015).

15. Scholefield, G., Whiting, R., Errington, J. & Murray, H. Spo0J regulates the oligomeric state of Soj to trigger its switch from an activator to an inhibitor of DNA replication initiation. Mol Microbiol 79, 1089–100 (2011).

16. Grilley, M., Welsh, K.M., Su, S.S. & Modrich, P. Isolation and characterization of the *Escherichia coli mutL* gene product. J Biol Chem 264, 1000–4 (1989).

17. Modrich, P. Methyl-directed DNA mismatch correction. J Biol Chem 264, 6597-600 (1989).

18. Sullivan, N.L., Marquis, K.A. & Rudner, D.Z. Recruitment of SMC by ParB-parS organizes the origin region and promotes efficient chromosome segregation. Cell 137, 697–707 (2009).

19. Wang, X., Montero Llopis, P. & Rudner, D.Z. *Bacillus subtilis* chromosome organization oscillates between two distinct patterns. Proc Natl Acad Sci U S A 111, 12877–82 (2014).

20. Hansen, F.G. & Atlung, T. The DnaA Tale. Front Microbiol 9, 319 (2018).

21. Leonard, A.C. & Grimwade, J.E. The orisome: structure and function. Front Microbiol 6, 545 (2015).

22. Ben-Yehuda, S. et al. Defining a centromere-like element in *Bacillus subtilis* by identifying the binding sites for the chromosome-anchoring protein RacA. Mol Cell 17, 773–82 (2005).

23. Ben-Yehuda, S., Rudner, D.Z. & Losick, R. RacA, a bacterial protein that anchors chromosomes to the cell poles. Science 299, 532–6 (2003).

24. Wang, X., Brandao, H.B., Le, T.B., Laub, M.T. & Rudner, D.Z. *Bacillus subtilis* SMC complexes juxtapose chromosome arms as they travel from origin to terminus. Science 355, 524–527 (2017).

25. Wu, L.J. & Errington, J. RacA and the Soj-Spo0J system combine to effect polar chromosome segregation in sporulating *Bacillus subtilis*. Mol Microbiol 49, 146–375 (2003).

26. Burton, B. & Dubnau, D. Membrane-associated DNA transport machines. Cold Spring Harb Perspect Biol 2, a000406 (2010).

27. Adams, D.W., Wu, L.J. & Errington, J. Nucleoid occlusion protein Noc recruits DNA to the bacterial cell membrane. EMBO J 34, 491–501 (2015).

28. Bernhardt, T.G. & de Boer, P.A. SlmA, a nucleoid-associated, FtsZ binding protein required for blocking septal ring assembly over chromosomes in *E*. coli. Mol Cell 18, 555–64 (2005).

29. Schumacher, M.A. & Zeng, W. Structures of the nucleoid occlusion protein SlmA bound to DNA and the C-terminal domain of the cytoskeletal protein FtsZ. Proc Natl Acad Sci U S A 113, 4988–93 (2016).

30. Cho, H., McManus, H.R., Dove, S.L. & Bernhardt, T.G. Nucleoid occlusion factor SlmA is a DNA-activated FtsZ polymerization antagonist. Proc Natl Acad Sci U S A 108, 3773–8 (2011).

31. Tonthat, N.K. et al. Molecular mechanism by which the nucleoid occlusion factor, SlmA, keeps cytokinesis in check. EMBO J 30, 154–64 (2011).

32. Rowlett, V.W. & Margolin, W. The Min system and other nucleoid-independent regulators of Z ring positioning. Front Microbiol 6, 478 (2015).

33. Bailey, M.W., Bisicchia, P., Warren, B.T., Sherratt, D.J. & Mannik, J. Evidence for divisome localization mechanisms independent of the Min system and SlmA in *Escherichia coli*. PLoS Genet 10, e1004504 (2014).

34. Buss, J.A., Peters, N.T., Xiao, J. & Bernhardt, T.G. ZapA and ZapB form an FtsZ-independent structure at midcell. Mol Microbiol 104, 652–663 (2017).

35. Wu, L.J. & Errington, J. Coordination of cell division and chromosome segregation by a nucleoid occlusion protein in *Bacillus subtilis*. Cell 117, 915–25 (2004).

36. Wu, L.J. et al. Noc protein binds to specific DNA sequences to coordinate cell division with chromosome segregation. EMBO J 28, 1940–52 (2009).

37. Webb, C.D. et al. Bipolar localization of the replication origin regions of chromosomes in vegetative and sporulating cells of *B*. subtilis. Cell 88, 667–74 (1997).

38. Wu, L.J. & Errington, J. Use of asymmetric cell division and *spoIIIE* mutants to probe chromosome orientation and organization in *Bacillus subtilis*. Mol Microbiol 27, 777–86 (1998).

39. Kloosterman, T.G. et al. Complex polar machinery required for proper chromosome segregation in vegetative and sporulating cells of *Bacillus subtilis*. Mol Microbiol 101, 333–50 (2016).

40. Gonzy-Treboul, G., Karmazyn-Campelli, C. & Stragier, P. Developmental regulation of transcription of the *Bacillus subtilis ftsAZ* operon. J Mol Biol 224, 967–79 (1992).

41. Levin, P.A. & Losick, R. Transcription factor Spo0A switches the localization of the cell division protein FtsZ from a medial to a bipolar pattern in *Bacillus subtilis*. Genes Dev 10, 478–88 (1996).

42. Ben-Yehuda, S. & Losick, R. Asymmetric cell division in *B. subtilis* involves a spiral-like intermediate of the cytokinetic protein FtsZ. Cell 109, 257–66 (2002).

43. Khvorova, A., Zhang, L., Higgins, M.L. & Piggot, P.J. The *spoIIE* locus is involved in the Spo0A-dependent switch in the location of FtsZ rings in *Bacillus subtilis*. J Bacteriol 180, 1256–60 (1998).

44. Frandsen, N., Barak, I., Karmazyn-Campelli, C. & Stragier, P. Transient gene asymmetry during sporulation and establishment of cell specificity in *Bacillus subtilis*. Genes Dev 13, 394–9 (1999).

45. Wang, S.T. et al. The forespore line of gene expression in *Bacillus subtilis*. J Mol Biol 358, 16–37 (2006).

46. Fiche, J.B. et al. Recruitment, assembly, and molecular architecture of the SpoIIIE DNA pump revealed by superresolution microscopy. PLoS Biol 11, e1001557 (2013).

47. Wu, L.J. & Errington, J. *Bacillus subtilis* SpoIIIE protein required for DNA segregation during asymmetric cell division. Science 264, 572–5 (1994).

48. Miller, A.K., Brown, E.E., Mercado, B.T. & Herman, J.K. A DNA-binding protein defines the precise region of chromosome capture during *Bacillus* sporulation. Mol Microbiol 99, 111–22 (2016).

49. Wagner-Herman, J.K. et al. RefZ facilitates the switch from medial to polar division during spore formation in *Bacillus subtilis*. J Bacteriol 194, 4608–18 (2012).

50. Britton, R.A. et al. Genome-wide analysis of the stationary-phase sigma factor (sigma-H) regulon of *Bacillus subtilis*. J Bacteriol 184, 4881–90 (2002).

51. Fujita, M., Gonzalez-Pastor, J.E. & Losick, R. High- and low-threshold genes in the Spo0A regulon of *Bacillus subtilis*. J Bacteriol 187, 1357–68 (2005).

52. Molle, V. et al. The Spo0A regulon of *Bacillus subtilis*. Mol Microbiol 50, 1683–701 (2003).

53. Gibson, D.G. et al. Enzymatic assembly of DNA molecules up to several hundred kilobases. Nat Methods 6, 343–5 (2009).

54. Yu, Z., Reichheld, S.E., Savchenko, A., Parkinson, J. & Davidson, A.R. A comprehensive analysis of structural and sequence conservation in the TetR family transcriptional regulators. J Mol Biol 400, 847–64 (2010).

55. Gibrat, J.F., Madej, T. & Bryant, S.H. Surprising similarities in structure comparison. Curr Opin Struct Biol 6, 377–85 (1996).

56. Agari, Y., Sakamoto, K., Kuramitsu, S. & Shinkai, A. Transcriptional repression mediated by a TetR family protein, PfmR, from Thermus thermophilus HB8. J Bacteriol 194, 4630–41 (2012).

57. Crowe, A.M. et al. Structural and functional characterization of a ketosteroid transcriptional regulator of Mycobacterium tuberculosis. J Biol Chem 290, 872–82 (2015).

58. Tonthat, N.K. et al. SlmA forms a higher-order structure on DNA that inhibits cytokinetic Z-ring formation over the nucleoid. Proc Natl Acad Sci U S A 110, 10586–91 (2013).

59. Schumacher, M.A. et al. Structural mechanisms of QacR induction and multidrug recognition. Science 294, 2158–63 (2001).

60. Grkovic, S., Brown, M.H., Schumacher, M.A., Brennan, R.G. & Skurray, R.A. The *Staphylococcal* QacR multidrug regulator binds a correctly spaced operator as a pair of dimers. J Bacteriol 183, 7102–9 (2001).

61. Cho, H. & Bernhardt, T.G. Identification of the SlmA active site responsible for blocking bacterial cytokinetic ring assembly over the chromosome. PLoS Genet 9, e1003304 (2013).

62. Cuthbertson, L. & Nodwell, J.R. The TetR family of regulators. Microbiol Mol Biol Rev 77, 440–75 (2013).

63. Engohang-Ndong, J. et al. EthR, a repressor of the TetR/CamR family implicated in ethionamide resistance in mycobacteria, octamerizes cooperatively on its operator. Mol Microbiol 51, 175–88 (2004).

64. Rodikova, E.A. et al. Two HlyIIR dimers bind to a long perfect inverted repeat in the operator of the hemolysin II gene from *Bacillus cereus*. FEBS Lett 581, 11906 (2007).

65. Singh, A.K. et al. Crystal Structure of Fad35R from *Mycobacterium tuberculosis* H37Rv in the Apo-State. PLoS One 10, e0124333 (2015).

66. Karimova, G., Pidoux, J., Ullmann, A. & Ladant, D. A bacterial two-hybrid system based on a reconstituted signal transduction pathway. Proc Natl Acad Sci U S A 95, 5752–6 (1998).

67. Barak, I. & Muchova, K. The positioning of the asymmetric septum during sporulation in *Bacillus subtilis*. PLoS One 13, e0201979 (2018).

68. Shiu-Hin Chan, D. et al. Structural insights into the EthR-DNA interaction using native mass spectrometry. Chem Commun (Camb) 53, 3527–3530 (2017).

69. Szwedziak, P., Wang, Q., Freund, S.M. & Lowe, J. FtsA forms actin-like protofilaments. EMBO J 31, 2249–60 (2012).

70. Choi, H., Min, K., Mikami, B., Yoon, H.J. & Lee, H.H. Structural and Biochemical Studies Reveal a Putative FtsZ Recognition Site on the Z-ring Stabilizer ZapD. Mol Cells 39, 814–820 (2016).

71. Schumacher, M.A., Huang, K.H., Zeng, W. & Janakiraman, A. Structure of the Z Ring-associated Protein, ZapD, Bound to the C-terminal Domain of the Tubulin-like Protein, FtsZ, Suggests Mechanism of Z Ring Stabilization through FtsZ Cross-linking. J Biol Chem 292, 3740–3750 (2017).

72. Ghosal, D., Trambaiolo, D., Amos, L.A. & Lowe, J. MinCD cell division proteins form alternating copolymeric cytomotive filaments. Nat Commun 5, 5341 (2014).

73. Zhang, H.N. et al. Cyclic di-GMP regulates *Mycobacterium tuberculosis* resistance to ethionamide. Sci Rep 7, 5860 (2017).

74. Harwood, C.R. & Cutting, S.M. Molecular biological methods for Bacillus, (Wiley, New York, NY, 1990).

75. Zhang, X.Z. & Zhang, Y. Simple, fast and high-efficiency transformation system for directed evolution of cellulase in *Bacillus subtilis*. Microb Biotechnol 4, 98–105 (2011).

76. Schneider, C.A., Rasband, W.S. & Eliceiri, K.W. NIH Image to ImageJ: 25 years of image analysis. Nat Methods 9, 671–5 (2012).

77. Otwinowski, Z. & Minor, W. [20] Processing of X-ray diffraction data collected in oscillation mode. Methods Enzymol 276, 307–326 (1997).

78. Sheldrick, G.M. A short history of SHELX. Acta Crystallogr A 64, 112–22 (2008).

79. Zwart, P.H. Anomalous signal indicators in protein crystallography. Acta Crystallographica Section D, Biological Crystallography 61, 1437–48 (2005).

80. Winn, M.D. et al. Overview of the CCP4 suite and current developments. Acta Crystallographica Section D, Biological Crystallography 67, 235–42 (2011).

81. Adams, P.D. et al. PHENIX: a comprehensive Python-based system for macromolecular structure solution. Acta Crystallographica Section D, Biological Crystallography 66, 213–21 (2010).

82. Cowtan, K. The Buccaneer software for automated model building. 1. Tracing protein chains. Acta Crystallographica Section D, Biological Crystallography 62, 1002–11 (2006).

83. Emsley, P., Lohkamp, B., Scott, W.G. & Cowtan, K. Features and development of Coot. Acta Crystallographica Section D, Biological Crystallography 66, 486–501 (2010).

## Supporting Information References

1. Wagner-Herman, J.K. et al. RefZ facilitates the switch from medial to polar division during spore formation in *Bacillus subtilis*. J Bacteriol 194, 4608–18 (2012).

2. Miller, A.K., Brown, E.E., Mercado, B.T. & Herman, J.K. A DNA-binding protein defines the precise region of chromosome capture during *Bacillus* sporulation. Mol Microbiol 99, 111–22 (2016).

3. Gibson, D.G. et al. Enzymatic assembly of DNA molecules up to several hundred kilobases. Nat Methods 6, 343–5 (2009).

4. Schumacher, M.A. et al. Structural mechanisms of QacR induction and multidrug recognition. Science 294, 2158–63 (2001).

5. Schumacher, M.A. et al. Structural basis for cooperative DNA binding by two dimers of the multidrug-binding protein QacR. EMBO J 21, 1210–8 (2002).

